# Exercise Conditioning Enhances Insulin Sensitivity with Metabolic and Glycemic Control Driven by Hepatic Silencing of HMGB1 in Type 2 Diabetic Mice

**DOI:** 10.1101/2025.04.27.650052

**Authors:** Gabriela Martinez Bravo, Prabu Paramasivam, Quiteria Jacquez, Gabriella Sandoval, Hannah Beason, Esmeralda Batista Ramirez, Jinhua Chi, Haiwei Gu, Flavio de Castro Magalhaes, Roberto Ivan Mota Alvidrez

**Author notes:** Correspondence: Roberto Ivan Mota Alvidrez, MD, MS) Assistant Professor, Pharmaceutical Sciences, College of Pharmacy, University of New Mexico, Research Incubator Building (RIB), 2703 Frontier Ave NE, Albuquerque, NM 87106+ Office: RIB room 299, Lab: RIB room 250, Preferred / Institutional.

## Abstract

Type 2 Diabetes (T2D) is characterized by chronic insulin resistance and inflammation. Exercise induces hyperglycemia control in T2D more than pharmacologic therapies alone. High Mobility Group Box 1 (HMGB1) is a ubiquitous nuclear factor for T2D progression. We hypothesize that hepatic silencing of HMGB1 in T2D and exercise will increase insulin sensitivity and shift metabolic routes, decrease liver damage and improve metabolic health in T2D mice.

**Methods:** Liver HMGB1 KO (HMGB1^Δ^) was induced with AAV8-TBG in HMGB1 flox mice (AAV8-EGFP was used as wildtype controls). T2D mice were induced with a low-dose STZ and a high-fat diet. T2D mice ran for four weeks (1hr daily). Dual Energy X-Ray Absorptiometry (DEXA), A1C, glycemia, HMGB1, metabolomics and metabolic cage analysis were performed pre- and post-exercise. RT-PCR, immunoblotting and immunohistochemistry for glucose metabolism markers were performed in tissue samples.

**Results:** T2D HMGB1^Δ^ mice show 20% less hyperglycemia compared to controls with a strong correlation between reduced HMGB1 levels and lower A1C following exercise. Additionally, fat percentage, liver damage, and glycogen storage were significantly reduced. These findings highlight the high potential effect of exercise potentiated has via HMGB1 governed mechanisms and highlights HMGB1 as a therapeutic target for managing hyperglycemia in T2D.

**ARTICLE HIGHLIGHTS:** HMGB1 hepatic silencing in Type 2 Diabetic mice is a promising therapeutic target for enhancing Insulin sensitivity. Combination of hepatic HMGB1 silencing and exercise potentiates the metabolic benefits and liver protection in Type 2 Diabetic mice. Shift in metabolic routes and carbohydrate metabolism due to regulation of hepatic HMGB1 expression makes HMGB1 an ideal target for pharmacologic intervention. However, our study also underscores the value of exercise as the primary focus of diabetic glucose control aided by enhanced therapeutic targeting of HMGB1 for chronic inflammation and hyperglycemia.

## 1. INTRODUCTION

Type 2 diabetes (T2D) is a metabolic disorder characterized by insulin resistance, chronic hyperglycemia, and systemic inflammation that affects millions of people worldwide(1). Chronic hyperglycemia in T2D leads to a wide array of complications, including cardiovascular disease, neuropathy, nephropathy, and non-alcoholic fatty liver disease (NAFLD)(2, 3). The liver is a central organ in T2D, modulates gluconeogenesis and glycogenolysis(4). Insulin resistance dysregulates these processes in T2D, perpetuated by chronic inflammation and hyperglycemia(5).

T2D inflammatory mediators significantly contribute to the progression of metabolic end-organ damage(1, 6). High mobility group box 1 (HMGB1) is pivotal in driving inflammation and exacerbating insulin resistance in T2D(7). Acetylated HMGB1 translocates to the cytoplasm and secreted as a damage-associated molecular pattern (DAMP). HMGB1 promotes inflammation activating the Receptor for Advanced Glycation End-products (RAGE) and Toll-Like Receptor 4 (TLR4). This activates Nuclear Factor kappa-light-chain-enhancer of activated B cells (NF-κB), Mitogen-Activated Protein Kinase (MAPK), and c-Jun N-terminal Kinase (JNK) pathways, leading to chronic inflammation(8, 9). Hyperglycemia and oxidative stress enhance HMGB1 acetylation/secretion amplifying hepatic inflammation(6, 10). Targeting HMGB1 represents a promising therapeutic approach to reduce inflammation and mitigate liver damage in T2D(11).

Lifestyle interventions, including regular physical activity improve metabolic health and reduce the risk of complications in individuals with T2D(12). Exercise exerts multiple beneficial effects, including enhancing insulin sensitivity and decreasing systemic inflammation(13). Studies have shown that exercise can also decrease the release of inflammatory factors like HMGB1, improving T2D(14, 15). The combined effects of exercise and HMGB1 knockdown have yet to be studied in improving metabolic health. An inducible HMGB1 knockout model shows 34% reduction of hyperglycemia in Streptozotocin-induced diabetic mice(6). In this study, we hypothesize that hepatic HMGB1 silenced T2D mice undergoing exercise training will evidence increase in systemic insulin sensitivity (improved glycemic control) and positive shift in energy and carbohydrate metabolism with liver damage protection in T2D.

## 2. MATERIALS AND METHODS

### 2.1. Animal model

HMGB1 Flox mice on BL/6 background were bred in-house at the UNM Animal Resource Facility. All experiments followed UNM regulations and the Institutional Animal Care and Use Committee guidelines. [Animal Welfare Assurance # D16-00228 (A3350-01), USDA Registration # 85-R-0014, protocol number 23-201405-HSC].

### 2.2. Liver Knockdown and Type 2 diabetes induction

6-week-old mice were intraperitoneally injected with either Adenovirus (AAV8) Tiroxin Binding Globulin-EGFP for control/ Wild Type (WT) or with AAV8-TBG iCre for liver knockdown (^Δ^) (Vector Bio Labs). After 2 weeks of induction of liver knockdown (HMGB1^Δ^), we began the T2D model. For T2D, mice were placed on a high-fat diet (Research Diets D12492/Rodent diet with 60 kcal% fat) for the remainder of the study, and streptozotocin (25 mg/kg) was injected intraperitoneally for 5 days. Mice were given 6 additional weeks to develop T2D(6). Starting at 10 weeks old, weights and glucose (glycemia was quantified with a commercial glucometer via submandibular blood puncture) were measured biweekly for all groups until the end of the study.

Fasting blood glucose levels were evaluated, and values above 300 mg/dL indicated type 2 diabetes(16). For non-T2D, mice were placed on normal chow for the remainder of the study to have them as a control.

### 2.3. Groups/housing

Mice were housed in standard rodent cages under a 12:12 light cycle and normal temperature conditions with ad libitum access to food and water. Mice were randomly divided into 8 groups: 1. Wild Type (WT) sedentary “healthy” (Non-T2D sedentary), 2. WT + exercise “healthy” (Non-T2D exercise), 3. WT sedentary type 2 diabetic (T2D sedentary), 4. WT + exercise T2D (T2D exercise), 5. HMGB1^Δ^ sedentary “healthy” (HMGB1^Δ^ non-T2D sedentary), 6. HMGB1^Δ^ + exercise “healthy” (HMGB1^Δ^ non-T2D exercise), 7. HMGB1^Δ^ sedentary T2D (HMGB1^Δ^ T2D sedentary), and 8. HMGB1^Δ^ + exercise T2D (HMGB1^Δ^ T2D exercise).

### 2.4. Exercise training in vivo

At around 14 weeks of age, a baseline DEXA Scan was performed on all mice groups. Then, exercise mice were acclimated to the treadmill (COLUMBUS INSTRUMENTS TREADMILL SIMPLEX II) environment and exercise routine for 1 week. Day 1 was exploratory, continued by slow walking for 10 minutes on day 2, to increase time and speed daily until day 5 (30 minutes at 60-70 mts/min). From 15 weeks to 18 weeks old, exercise groups were placed on the treadmill at 80 mts/min, 1 hour/day, 5 days/week (4 weeks total). Sedentary mice were not exercised. At around 18 weeks old, post-exercise and sedentary mice underwent DEXA Scan measurements. After DEXA Scan, mice were moved to metabolic cages for 3 days.

### 2.5. Oral glucose tolerance test (OGTT)

OGTT was performed at baseline and post-exercise in T2D and HMGB1^Δ^ T2D mice that underwent a 16-h fasting period. Subsequently, the mice were given an oral glucose solution at 2 grams per kilogram of body weight. Blood glucose levels were then measured at 0, 30, 60, 90, and 120 minutes after the glucose administration. A glucometer was used to assess glucose tolerance(17).

### 2.6. Euthanasia; tissue collection and preparation

At 19 weeks of age, mice were anesthetized using isoflurane and humanely euthanized via cardiac puncture exsanguination while collecting whole blood. Liver, skeletal muscle, and adipose tissue were harvested for further analysis. Tissue collection and preparation as well as liver function assessment was performed as previously described [6].

### 2.7. Liver and plasma untargeted metabolomic

The untargeted LC-MS metabolomics methods and analysis were performed from prior established studies (18, 19). Reagents used in this study included acetonitrile (ACN), methanol (MeOH), ammonium acetate, and acetic acid, all LC-MS grade and sourced from Fisher Scientific (Pittsburgh, PA). Ammonium hydroxide was obtained from Sigma-Aldrich (Saint Louis, MO), while deionized water (DI water) was supplied a Water Purification System from EMD Millipore (Billerica, MA). Phosphate-buffered saline (PBS) was purchased from GE Healthcare Life Sciences (Logan, UT). Standard compounds corresponding to the measured metabolites were acquired from Sigma-Aldrich (Saint Louis, MO) and Fisher Scientific (Pittsburgh, PA).

### 2.8. Statistical analysis

CalR software was used to analyze data retrieved from the metabolic cages, then plotted using GraphPad Prism v8.4. Metaboanalyst online software was implemented for metabolic analysis to retrieve the graphs used in this research. All illustrated data was performed using GraphPad Prism v8.4 (GraphPad Software, Inc., USA). Unpaired two-tailed Student’s t-tests and one-way ANOVA were performed when appropriate. Analyses are detailed in each section and each figure legend. Figure legends specify the number of replicates and samples analyzed. P-values <0.05 were determined to be significant. Data are presented as mean ± standard error (SE) unless otherwise indicated. The specific number of animals is detailed for each experiment/assay for each figure/procedure. Overall, an n of 3-8 mice were used per group per experiment. Statistical significance was set at *: <0.05, **: 0.001–0.01, ***: 0.0001–0.001, and ****: <0.0001.

## 3. RESULTS

### 3.1. Hepatic HMGB1 Silencing Reduces HMGB1 Expression and Secretion, with decreasing Liver Damage in T2D Mice

HMGB1 qRT-PCR and protein analyses demonstrate a significant reduction in HMGB1 gene/protein levels in isolated hepatocytes of HMGB1^Δ^ non-T2D and HMGB1^Δ^ T2D mice (Fig.1A), compared to their non-silenced counterparts (Fig. 1B and 1C). Circulating HMGB1 is 50% lower in HMGB1^Δ^ T2D mice (Fig. 1D). Aspartate transaminase (AST) was reduced in HMGB1^Δ^ T2D mice (Fig. 1E). Histological analysis correlated these findings with reduced lipid accumulation and inflammatory infiltrates compared to T2D mice with HMGB1 wildtype T2D mice (Fig.1F).

**Figure 1.**
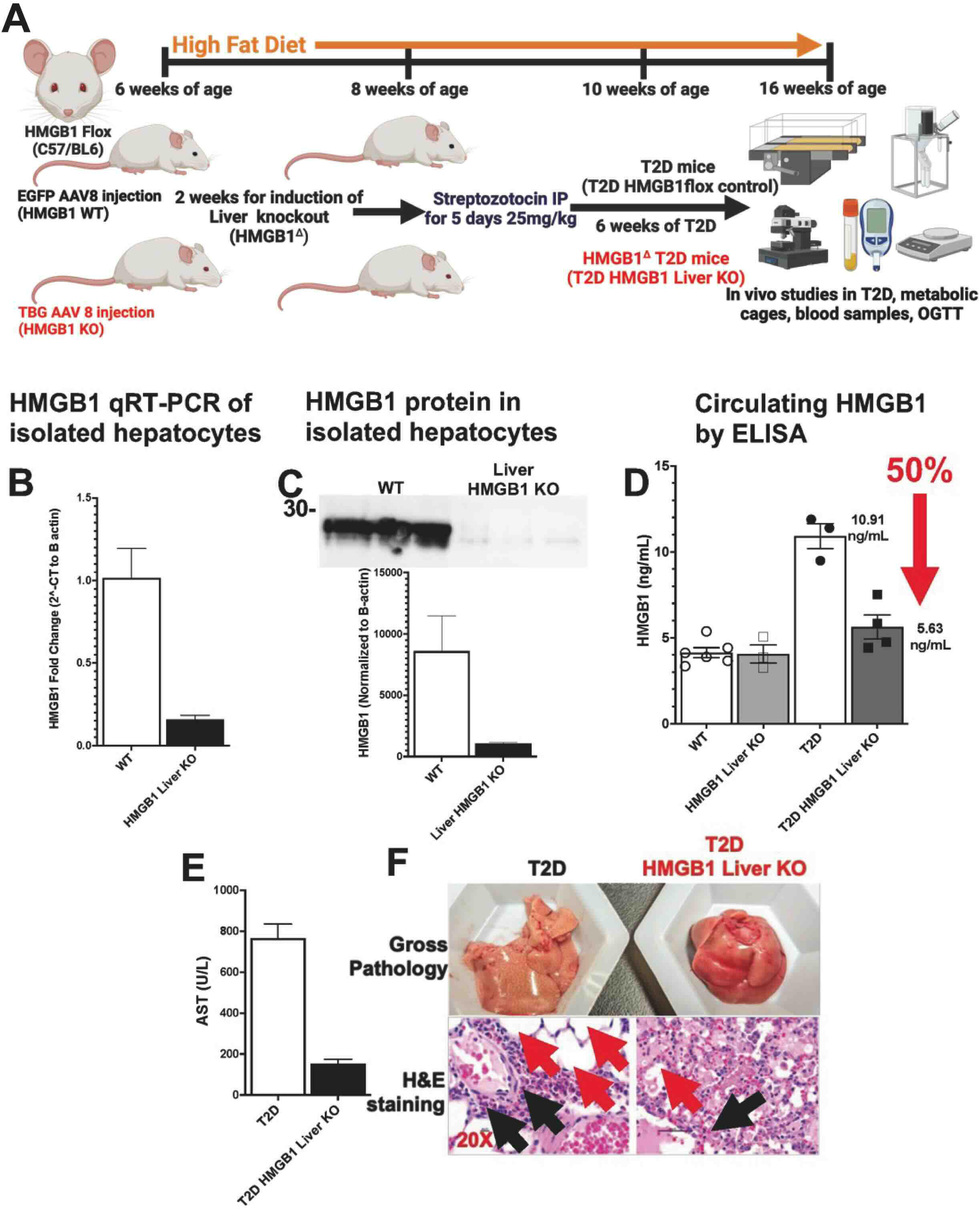
Effective silencing of hepatic HMGB1 expression and secretion in a T2D mouse model particularly in sedentary non-T2D vs T2D isolated hepatocytes. A) Flow diagram of the methodology for the in vivo silencing of hepatic HMGB1 using AAV8-TBG directed injections in HMGB1 flox mice in the T2D mouse model, B) qRT-PCR analysis of decreased HMGB1 gene expression in isolated hepatocytes from HMGB1Δ non-T2D (n=3) mice vs non-T2D (n=3), C) Representative Western blot showing the decrease of HMGB1 protein levels in HMGB1Δ non-T2D (n=3) vs non-T2D (n=3) mice hepatocytes (bar graph summarizing the relative levels of HMGB1), D) ELISA quantification of circulating HMGB1 from non-T2D (n=5), HMGB1Δ non-T2D (n=3), T2D (n=3) and HMGB1Δ T2D mice (n=4), E) Bar graph summarizing results from Aspartate transaminase (AST) in plasma showing decreased liver damage in HMGB1Δ T2D mice (n=4) compared to T2D mice (n=4), F) Histological analysis shows increase lipid accumulation and pale aspect of T2D mice compared to HMGB1Δ T2D mice in macroscopic analysis post tissue harvesting; cryosections of liver H&E stained slides of liver in T2D and HMGB1Δ T2D from mice (Black arrows point to inflammatory cellular infiltrates, red arrows point to lipid droplets). All data is represented as mean±SEM. All non-T2D, HMGB1Δ non-T2D, T2D and HMGB1Δ T2D are used for One-way ANOVA analysis. Significance levels: *:<0.05, **: 0.001-0.01, ***: 0.0001, ****: <0.0001.

### 3.2. Exercise Improves Glycemic Control by Decreasing Glycemia, A1C, and HMGB1 Levels in HMGB1^Δ^ T2D Mice

Exercise in hepatic HMGB1 silenced T2D mice demonstrated improved glycemia, A1C levels, and circulating HMGB, especially in HMGB1^Δ^ T2D mice (Fig. 2A). HMGB1^Δ^ T2D mice have higher glucose tolerance due to hepatic HMGB1 silencing. Post-training HMGB1^Δ^ T2D mice show enhanced glucose tolerance compared to sedentary WT T2D mice (Fig. 2B). HMGB1^Δ^ T2D mice fasting glucose levels decreased by 31.13% from baseline to the end of the study. In contrast, regular T2D mice exhibited a minor reduction of 12% in fasting glucose levels over the same timeframe of the study (Fig. 2C). Exercise reduces circulating HMGB1 levels in HMGB1^Δ^ T2D even though it was not statistically significant, however a difference was observed post exercise between HMGB1^Δ^ non-T2D and HMGB1^Δ^ T2D (p = 0.0008). Contrarily to T2D control mice, that show a slight increase in HMGB1, HMGB1 liver silencing in T2D aids in decreasing circulating HMGB1 post exercise (Fig. 2D). A1C levels show a decreased post-exercise in HMGB1^Δ^ T2D mice compared to sedentary T2D (p<0.0001)(Fig. 2E). Mouse weight decreased during the first two weeks in HMGB1^Δ^ T2D group, as expected, as they adapted to the exercise regimen, weight increased (Fig.2F).

**Figure 2.**
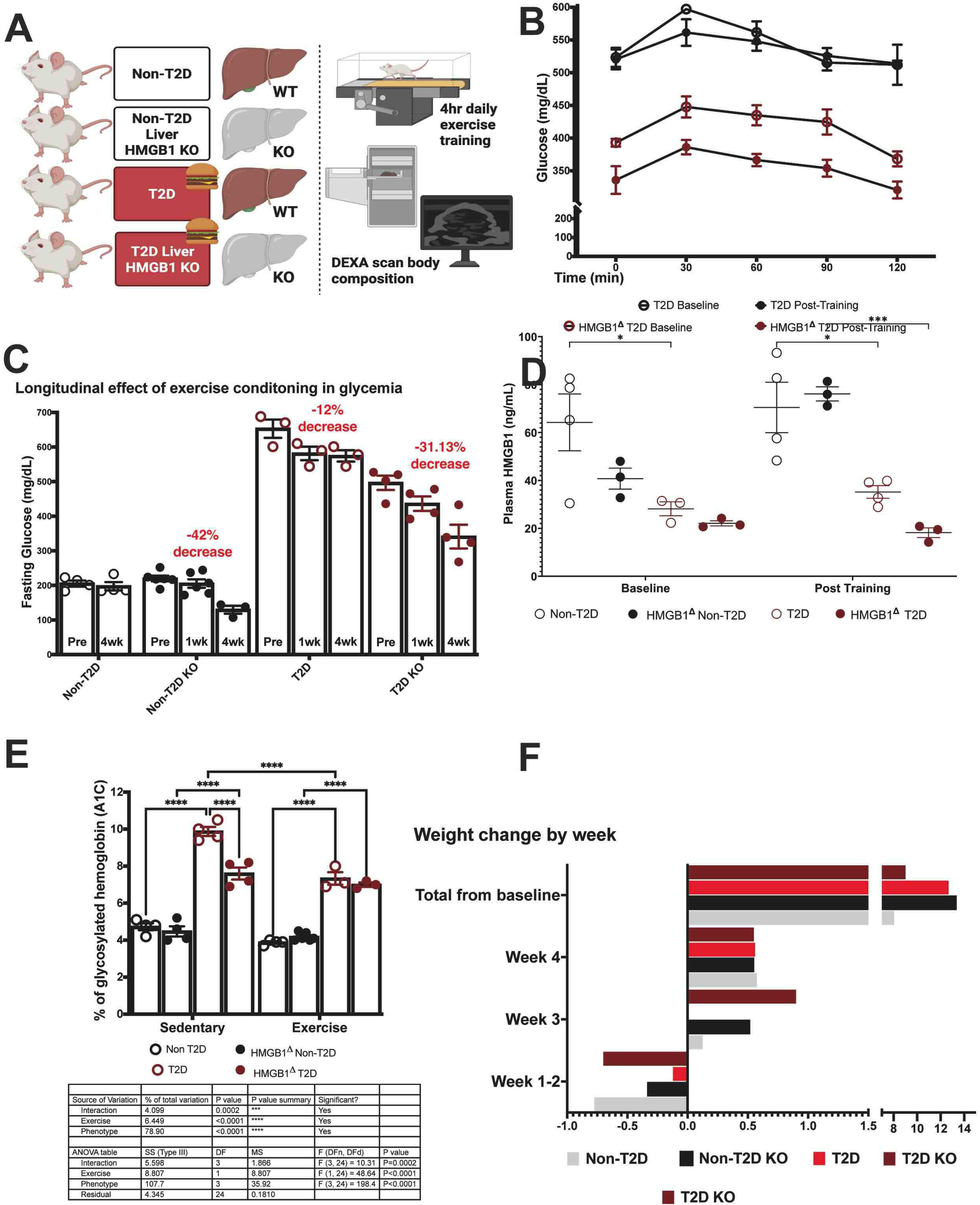
Glucose, A1C and HMGB1 levels share associated and potentially causal decreases post-training in HMGB1^Δ^ T2D mice. A) Diagram of the different mouse models used in this study, followed by the procedure performed, B) Glucose oral tolerance test (OGTT) shows glucose sensitivity through 120 minutes comparing T2D (n=5) and HMGB1^Δ^ T2D (n=5) mice at baseline (pre-training) and post-training, C) Bar graph summarizing the longitudinal effect of exercise condition in glycemia particularly in fasting glucose levels of non-T2D (n=4), HMGB1^Δ^ non-T2D (n=6), T2D (n=3) and HMGB1^Δ^ T2D (n=4) mice from baseline (pre-training), 1 week of training and 4 weeks of training (post-training), D) ELISA of circulating HMGB1 quantification from non-T2D (n=4), HMGB1^Δ^ non-T2D (n=3), T2D (n=3) and HMGB1^Δ^ T2D mice (n=3) from baseline (pre-training) and post-training, E) Bar graph summarizing the percentage levels of non-T2D (n=4), HMGB1^Δ^ non-T2D (n=4), T2D (n=3) and HMGB1^Δ^ T2D mice (n=3) from sedentary vs exercise mice with ANOVA table included, F) Bar graph showing the changes in weight by week from non-T2D (n=4), HMGB1^Δ^ non-T2D (n=4), T2D (n=3) and HMGB1^Δ^ T2D mice. All data is represented as mean±SEM. All non-T2D, HMGB1^Δ^ non-T2D, T2D, and HMGB1^Δ^ T2D are used for One-way and two way-ANOVA analysis. Significance levels: *:<0.05, **: 0.001-0.01, ***: 0.0001, ****: <0.0001.

### 3.3. Exercise Training Enhances Insulin Pathway Activation and Metabolites in HMGB1^Δ^ T2D Mice

We found notable fold changes in liver metabolites (Fig. 3A) with distinct metabolic responses to exercise in diabetic condition (Fig. 3B). Partial Least Squares Discriminant Analysis (PLS-DA) plot shows the variable importance in projection (VIP), in third position there is the 2-(beta-D-Glucose) that decrease in HMGB1^Δ^ T2D (Fig. 3C) and Random Forest plot, features ranked metabolites by their contributions to classification accuracy (Mean Decrease Accuracy) in first position there is L-Tyrosine that increase in HMGB1^Δ^ T2D, further illustrated these metabolic shifts, indicating a clear separation between the two groups based on liver metabolite profiles (Fig. 3D). QRT-PCR analysis revealed increased expression of insulin signaling genes, including Protein Kinase B (AKT) and Insulin Receptor Substrate (INRS), and upregulation of metabolic regulators such as Nuclear Factor Erythroid 2–Related Factor 2 (NRF2) and Forkhead Box O1(FOX01) in exercised HMGB1^Δ^ T2D mice (Fig. 3E). Immunoblot analysis further confirmed higher levels of phosphorylated AKT (p-AKT) in liver of the exercised group (Fig. 3F-G).

**Figure 3.**
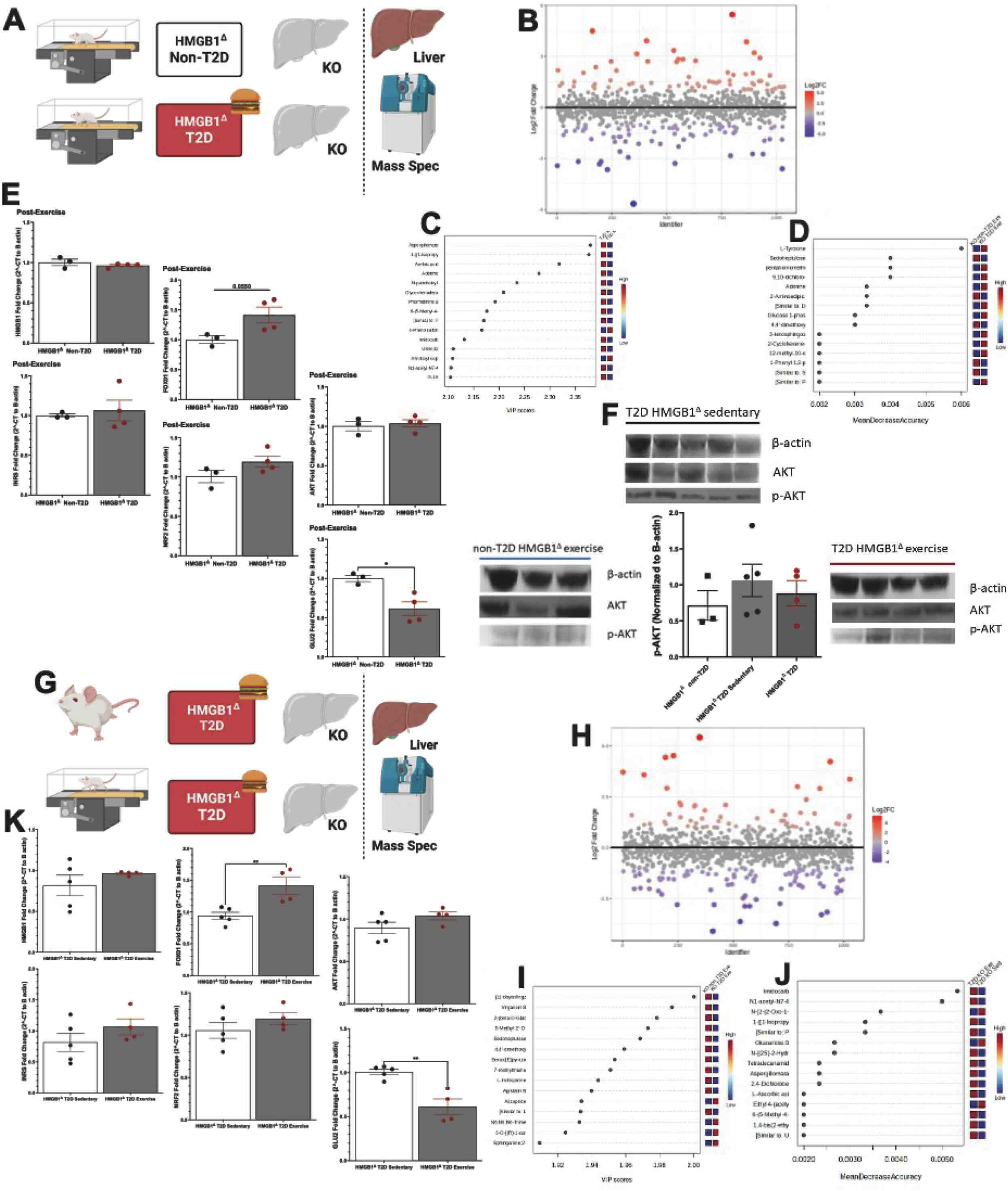
Exercise enhances Insulin pathway activation and post-training metabolite cumulates in liver from HMGB1^Δ^ T2D. A) Diagram of mouse models used in this study post-exercise HMGB1^Δ^ non-T2D and HMGB1^Δ^ T2D, followed by the procedure performed. B) Fold change (FC) in metabolites levels in liver tissues, C)Partial Least Squares Discriminant Analysis (PLS-DA), D) Random Forest plot, E) Quantitative PCR analysis of HMGB1, FOX01, AKT, INRS, NRF2 and GLU2 gene expression in hepatocytes from HMGB1^Δ^ non-T2D and HMGB1^Δ^ T2D mice, F) Representative Western blot showing AKT, p-AKT and HMGB1 protein levels in hepatocytes from HMGB1^Δ^ non-T2D and HMGB1^Δ^ T2D mice, G) Diagram of mouse models used in this study pre-exercise HMGB1^Δ^ T2D and post-exercise HMGB1^Δ^ T2D, followed by the procedure performed. H) Fold change (FC) in metabolites levels in liver tissues, I)Partial Least Squares Discriminant Analysis (PLS-DA), J) Random Forest plot, K) Quantitative PCR analysis of HMGB1, FOX01, AKT, INRS, NRF2, and GLU2 gene expression in hepatocytes from pre-exercise HMGB1^Δ^ T2D and post-exercise HMGB1^Δ^ T2D mice. All data represented mean±SEM. N of 3 for HMGB1^Δ^ non-T2D exercise, 4 for HMGB1^Δ^ T2D exercise, and 5 for HMGB1^Δ^ T2D sedentary. All data are used for One-way ANOVA analysis. Significance levels: *:<0.05, **: 0.001-0.01, ***: 0.0001, ****: <0.0001.

Exercise induces significant shifts in metabolic pathways (Fig. 3H). We found differential accumulation of metabolites associated with energy metabolism and glucose homeostasis (Fig. 3I, 3J). RT-Q PCR analysis shows upregulation in insulin pathway-related genes in the post-exercise group (Fig. 3K). Post-exercise HMGB1^Δ^ T2D mice demonstrated significant changes in the abundance of glucose metabolites and the different pathways activated in supplementary figure 2, compared to pre-exercise controls.

### 3.4. Metabolic Cage Analysis Evidence Enhanced Oxygen Consumption and Carbohydrate Metabolic Efficiency Driven by Exercise in HMGB1^Δ^ T2D Mice

Exercise improves metabolic parameters, including oxygen consumption, energy expenditure, and carbohydrate metabolism in HMGB1^Δ^ T2D mice compared to non-T2D and T2D controls (Fig. 4A). Metabolic cage analysis shows exercise in HMGB1^Δ^ T2D mice increase oxygen consumption (Fig. 4C) and carbon dioxide production (Fig. 4D). We also evidenced how energy expenditure (Fig. 4E) and respiratory exchange ratio (Fig. 4F) improved systemic glycolytic capacity and improved carbohydrate metabolism in HMGB1^Δ^ T2D mice undergoing exercise.

**Figure 4.**
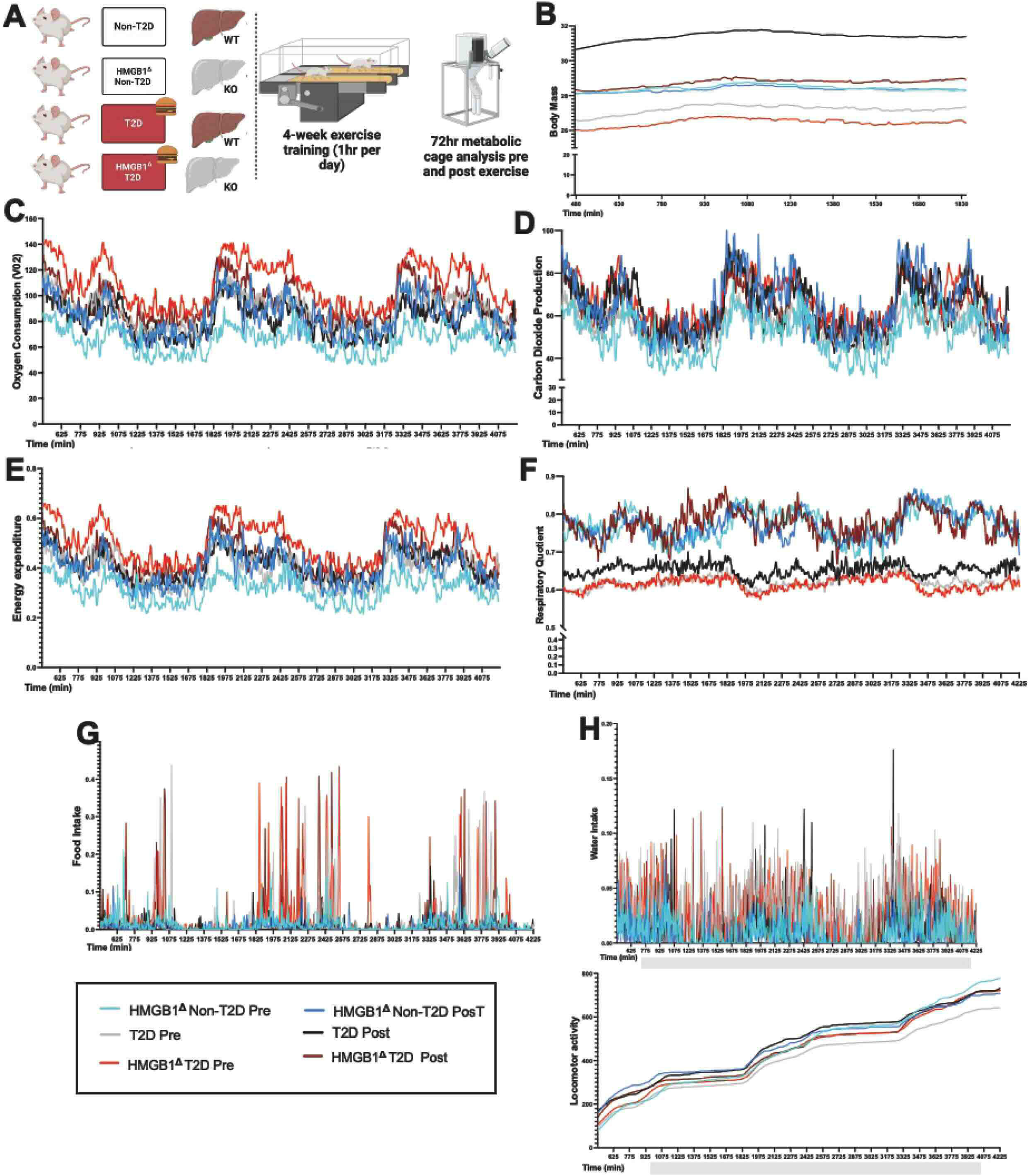
Metabolic cage analysis evidenced enhanced oxygen consumption and carbohydrate metabolism shifts post-exercise conditioning in HMGB1^Δ^ T2D. A) Diagram of the different mouse models used in this study, followed by the procedure performed on non-T2D, HMGB1^Δ^ non-T2D, T2D, and HMGB1^Δ^ T2D mice, B) Linear graph representing the difference in body mass pre and post-exercise, C) Linear graph representing the difference in oxygen consumption pre and post-exercise, D) Linear graph representing the difference in Carbon dioxide production pre and post-exercise, E) Linear graph representing the difference in Energy Expenditure pre and post-exercise, F) Linear graph representing the difference in Respiratory rate pre and post-exercise, G) Linear graph representing the difference in Food intake pre and post-exercise, H) Linear graph representing the difference in Water Intake pre and post-exercise, I) Linear graph representing the difference in locomotor activity pre and post-exercise. All data is represented as mean±SEM. N of 4 for non-T2D, of 4 for HMGB1^Δ^ non-T2D, of 4 for T2D and of 4 for HMGB1^Δ^ T2D.

Changes in body mass (Fig. 4B), food intake (Fig. 4G), and water intake (Fig. 4H) post-exercise indicate a shift in metabolic balance post exercise in HMGB1^Δ^ T2D mice. Supplementary Figure 3 shows the enhanced effects in energy expenditure, oxygen consumption and respiratory quotient in age matched sedentary HMGB1^Δ^T2D and T2D mice.

### 3.5. Circulating Plasma Metabolites Reveal Distinct Metabolic Changes in HMGB1^Δ^ T2D Mice Pre- and Post-Exercise

Heat Map Analysis revealed significant clustering of metabolic changes in our groups (Fig. 5A). HMGB1^Δ^ non-T2D mice exhibited a distinct decreased metabolic profile compared to HMGB1^Δ^ T2D (Fig. 5B). The Fold Change plot shows the significant upregulated and downregulated metabolites in HMGB1^Δ^ T2D (Fig. 5C). VIP scores and SPLS plots confirmed metabolic differences between HMGB1^Δ^ T2D and non-T2D mice some of the notorious changes are DL-Carnitine, 3-formyl-l-tyr a derivative of the amino acid tyrosine, 3-formyl-l-tyr a derivative of the amino acid tyrosine, and 2-amino-3 methyl that increase post-exercise in HMGB1^Δ^ T2D; dimethylol prop that decreased post-exercise in HMGB1^Δ^ T2D (Fig. 5D, E, F). HMGB1^Δ^ T2D mice have lower Methylone and 2-butyne-1,4-diol post-exercise (Fig. 5I, J). Plasma metabolite heat map shows metabolic shifts induced by physical activity (Fig. 5G). Exercise improved key metabolic pathways disrupted by T2D (Fig. 5H). Supplementary Figure 1 identified clustered differences in metabolite profiles between pre-exercise T2D and non-T2D mice. Supplementary Figure 2 shows the pathways altered between non-T2D and T2D mice.

**Figure 5.**
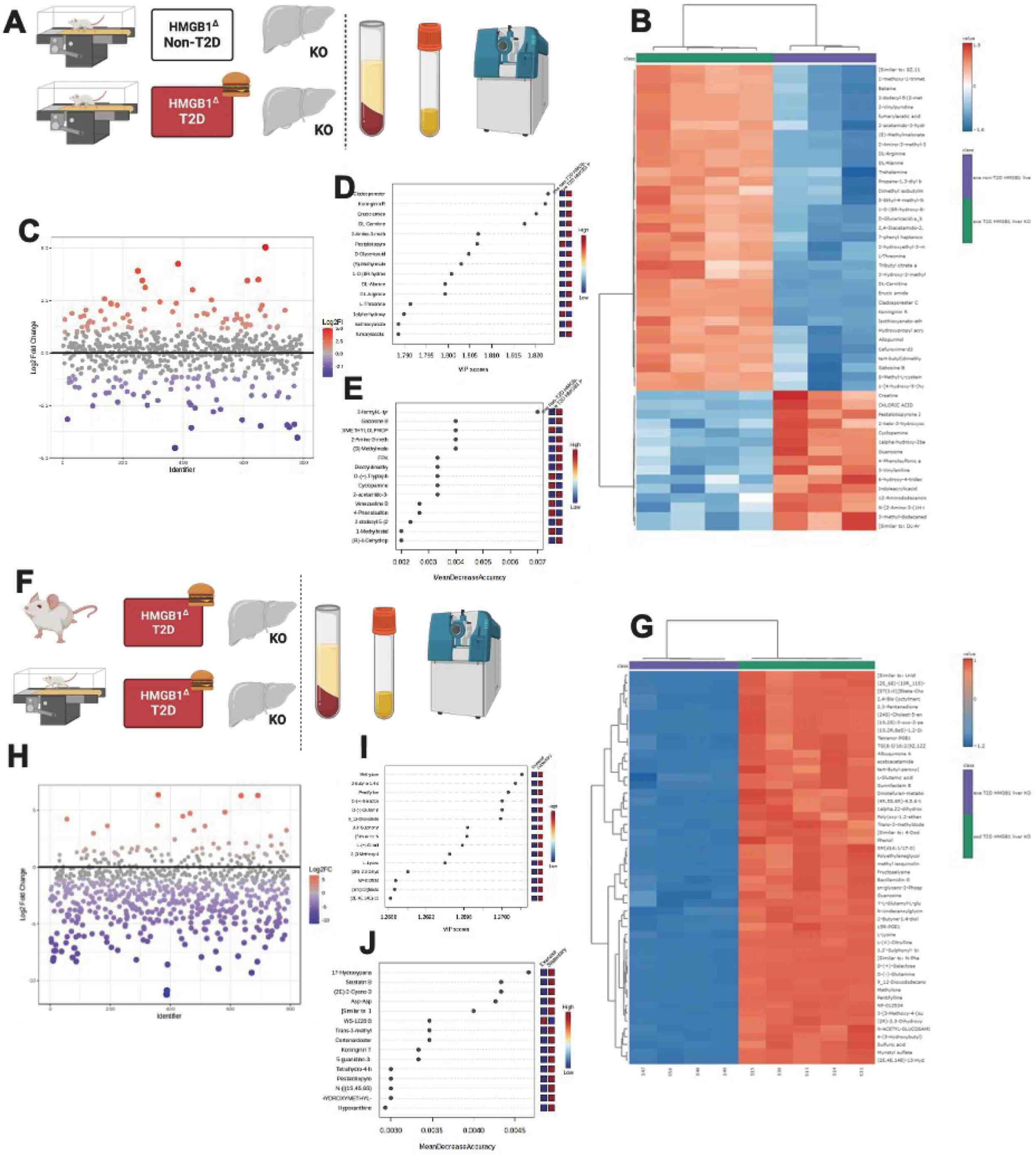
Plasma metabolomics identified overriding circulating metabolites in HMGB1^Δ^ T2D mice post-exercise. A) Diagram of the different mouse models used in this study post-exercise to evaluate the effect of T2D in hepatic HMGB1 silenced mice comparing HMGB1^Δ^ non-T2D and HMGB1^Δ^ T2D, followed by the procedure performed, B) Heat map displays the up- and down-regulated metabolites in the plasma post-exercise, C) Fold change plot of plasma metabolites in plasma samples, D) VIP score plots and E) SPLS plots of ranked metabolites differences in plasma samples, F) Diagram of mouse models used in this study to evaluate the effect of exercise comparing sedentary HMGB1^Δ^ T2D and post-exercise HMGB1^Δ^ T2D, followed by the procedure performed, G) Heat map displays the up- and down-regulated metabolites in plasma, H) Fold change plot of plasma metabolites in plasma samples comparing sedentary HMGB1^Δ^ T2D and post-exercise HMGB1^Δ^ T2D, I) VIP score plots and J) SPLS plots of ranked metabolites differences in plasma samples comparing sedentary HMGB1^Δ^ T2D and post-exercise HMGB1^Δ^ T2D. All data is represented as mean±SEM. N of 3 for HMGB1^Δ^ non-T2D exercise, 4 for HMGB1^Δ^ T2D exercise and 5 for HMGB1^Δ^ T2D sedentary.

### 3.6. Exercise Training Enhances Bone Mineral Density and Lean Muscle Mass While Reducing Body Fat in HMGB1^Δ^ T2D Mice

Exercise training improved bone mineral density, increased lean muscle mass, and reduced body fat percentage in HMGB1^Δ^ T2D mice (Fig. 6A, B). Bone mineral density significantly increases post-exercise in HMGB1^Δ^ T2D (p:0.0090) (Fig. 6C). Additionally, an increase in lean muscle mass in the exercise group was observed (Fig. 6D). Body fat percentage was decreased in both HMGB1^Δ^ T2D and HMGB1^Δ^ non-T2D mice after exercise (Fig. 6E, F).

**Figure 6.**
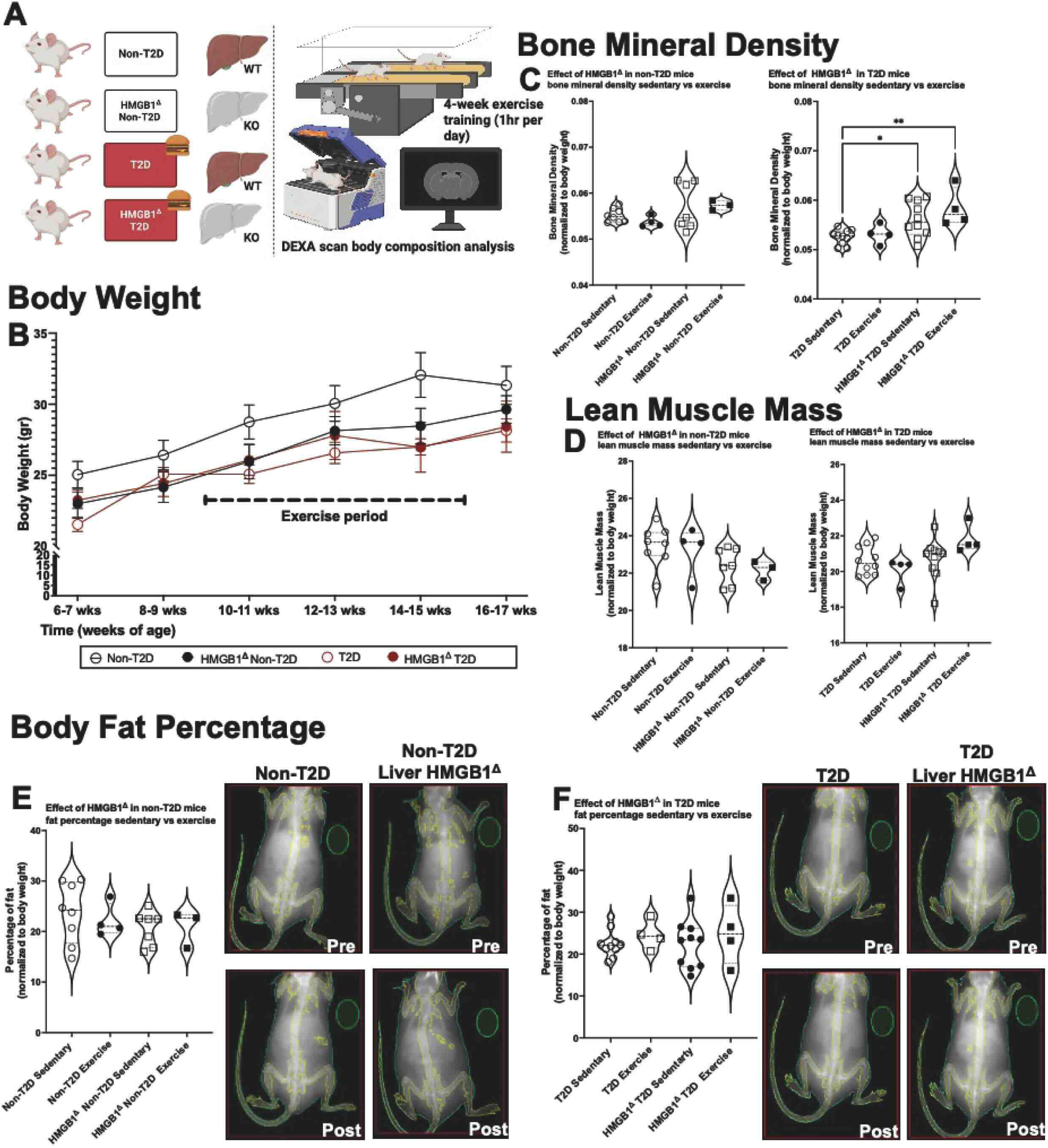
4-weeks of exercise conditioning training increases bone mineral density and lean muscle mass in HMGB1^Δ^ T2D mice. A) Diagram of the different mouse models used in this study, followed by the procedure performed in HMGB1^Δ^ T2D (n=7 for sedentary) (n=3 for exercise) and nonT2D (n=8 for sedentary) (n=4 for exercise) mice, B) Linear plot representing the change in weight during the exercise period, C) Violin plot representing the effect of in bone mineral density sedentary vs. exercise, D) Violin plot representing the effect in Lean Muscle Mass sedentary vs. exercise, E) Violin plot representing the effect of HMGB1^Δ^ non-T2D mice in body fat percentage sedentary vs. exercise, showing a decrease in the exercise cohort, F) Violin plot representing the effect of HMGB1Δ T2D mice in body fat percentage sedentary vs. exercise, showing a decrease in the exercise cohort. All data is represented as mean±SEM. All non-T2D, HMGB1^Δ^ non-T2D, T2D, and HMGB1^Δ^ T2D are used for One-way ANOVA analysis. Significance levels: *:<0.05, **: 0.001-0.01, ***: 0.0001, ****: <0.0001.

### 3.7. Exercise Modulates Metabolite Profile in Liver and Skeletal Muscle of HMGB1^Δ^ T2D Mice, Inhibits HMGB1 Release from Skeletal Muscle Cells

Venn diagrams show 21 metabolites significantly upregulated and 29 downregulated in the liver (Fig. 7A). In contrast, skeletal muscle showed only 12 upregulated and 9 downregulated metabolites (Fig. 7B). The volcano plot shows distinct patterns between liver and skeletal muscle with the liver displaying more extensive metabolite changes compared to skeletal muscle (Fig. 7C, D). In our other cohort, Venn diagrams show more considerable significant differences between the two groups of mice, with 34 metabolites being significantly upregulated and 19 downregulated in the liver, with skeletal muscle showing 12 upregulated and 32 downregulated metabolites (Fig. 7E-F). The volcano plot shows graphically these significant changes post-exercise in HMGB1^Δ^ T2D mice (Fig. 7G, H). Finally, in vitro results presented in Supplemental Figure 4 using electrical pulse stimulation (EPS) were performed to mimic skeletal muscle contraction in differentiated C2C12 skeletal muscle cells. Results show hoe HMGB1 release was significantly inhibited after EPS. Immunofluorescence imaging confirmed that HMGB1 localization was reduced in the cytoplasm/nucleus of stimulated cells. Additionally, qRT-PCR and Western blot analyses demonstrated that mimicking contractions in the cell reduced the cells’ HMGB1 gene expression in normal conditions and protein release in all conditions.

**Figure 7.**
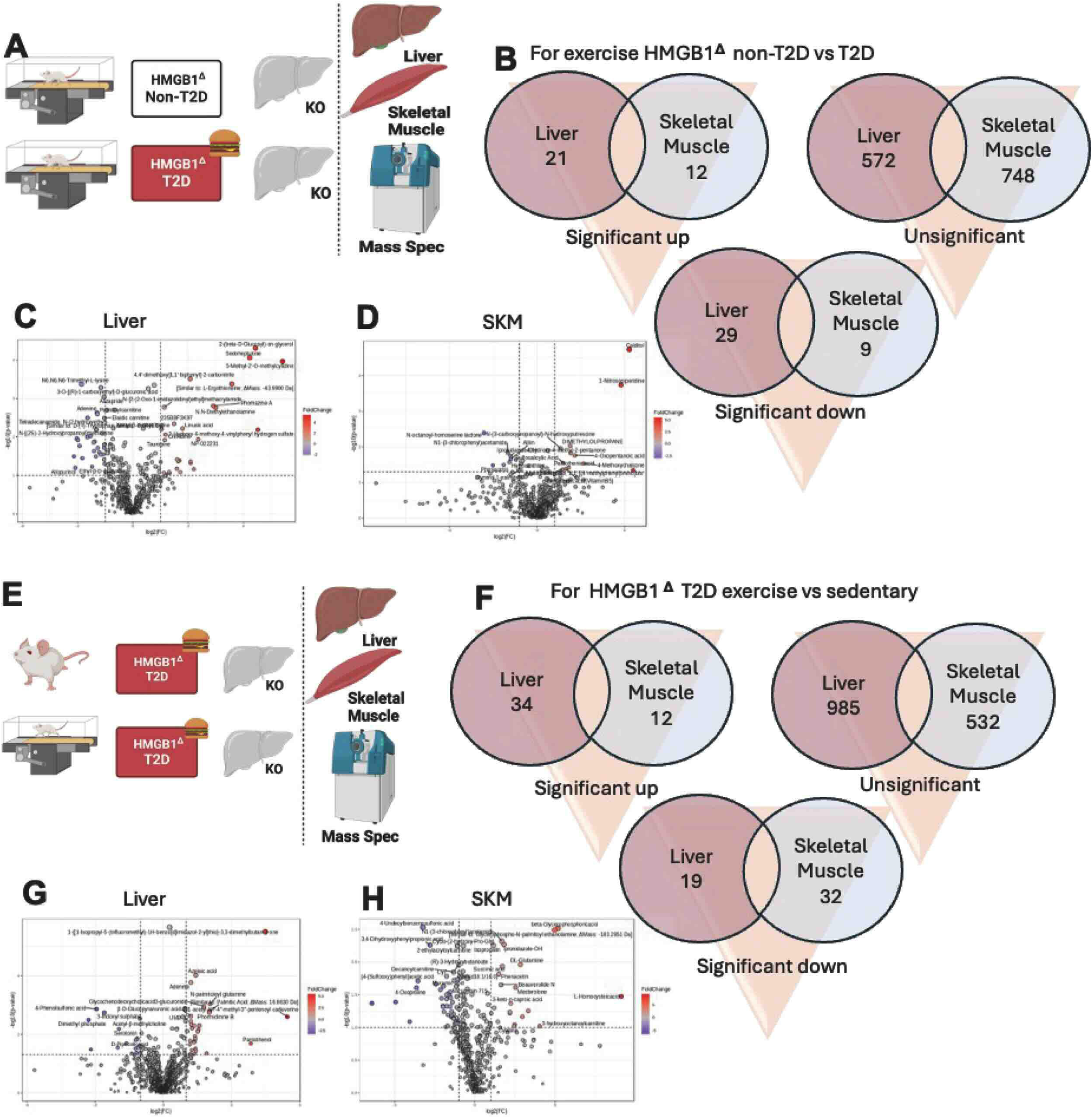
Volcano plot analysis identifies differences in significantly increased and decreased regulated metabolites in the liver and skeletal muscle of HMGB1^Δ^ T2D mice post-exercise. A) Diagram of the different mouse models used in this study post-exercise HMGB1^Δ^ non-T2D and HMGB1^Δ^ T2D, followed by the procedure performed, B) Differential findings of significant up, down, and insignificant metabolites present in the volcano plots from liver and skeletal muscle, C) Volcano plot (AB-Sciex QTOF) from the liver metabolites comparing the effect of T2D in hepatic HMGB1 silenced mice, D) Volcano plot from the skeletal muscle metabolites comparing the effect of T2D in hepatic HMGB1 silenced mice, E) Diagram of the different mouse models used in this study post-exercise HMGB1^Δ^ non-T2D and HMGB1^Δ^ T2D, followed by the procedure performed, F) Differential findings of significant up, down, and insignificant metabolites present in the volcano plots from liver and skeletal muscle, G) Volcano plot (AB-Sciex QTOF) from the liver metabolites comparing the effect of exercise in HMGB1^Δ^ T2D mice, H) Volcano plot from the skeletal muscles metabolites comparing the effect of exercise in HMGB1^Δ^ T2D mice. All data is represented as mean±SEM. N of 3 for HMGB1^Δ^ non-T2D exercise, 4 for HMGB1^Δ^ T2D exercise and 5 for HMGB1^Δ^ T2D sedentary.

## 4. DISCUSSION

Our group has revealed for the first time how silencing hepatic HMGB1 prevents liver damage and inflammation in T2D mice with enhanced liver metabolite profile and systemic metabolic shift toward glucose control and carbohydrate metabolism.

Importantly, exercise combined with HMGB1 silencing improves glycemic control, enhances carbohydrate metabolism, and reduces T2D-associated liver disease via activation of insulin signaling pathways. Specifically, we observed significant reductions in circulating HMGB1 levels in HMGB1^Δ^ T2D mice post-exercise, correlating with improved fasting glucose, A1C (Fig. 2), and Akt phosphorylation (Fig. 3), a critical step in insulin signaling(13). These results suggest that HMGB1 silencing enhances the Akt-FoxO1 pathway, suppressing FoxO1-driven transcription of gluconeogenic genes such as G6pc and Pck1, consistent with findings from Matsumoto et al. (20) and Puigserver et al.(21). FoxO1, which regulates gluconeogenesis and lipogenesis (22), was modulated with hepatic metabolites (Fig. 7). Furthermore, exercise induced improvements in skeletal muscle insulin signaling and metabolite profiles (Fig. 3F, 7H) support a systemic enhancement of energy metabolism. Our results highlight the interplay between HMGB1 silencing, Akt activation, and FoxO1 suppression, suggesting a mechanistic link that extends beyond hepatic effects to systemic metabolic improvements(20).

The decrease in circulating HMGB1 levels, evidenced by a 50% reduction in plasma HMGB1 in HMGB1^Δ^ T2D mice, underscores the role of hepatic HMGB1 in mediating liver inflammation and highlights the therapeutic potential of its silencing(23). This is a very novel finding that evidence that in T2D, HMGB1 is not only primarily released/expressed by immune cells, but almost half of all HMGB1 derives directly from hepatocytes due to our results in circulating and resident HMGB1 (from isolated hepatocytes). Post-translational modifications, such as acetylation, can regulate this release, which promotes its translocation from the nucleus to the extracellular space. From parallel studies and published literature, we have learned that proinflammatory HMGB1, acetyl-HMGB1, is implicated in driving chronic inflammation, making it a critical target for therapeutic interventions(24, 25). Another important highlight shows how HMGB1 silencing protects from liver inflammatory complications in T2D is the reduction of AST levels that indicate less liver injury (Fig. 1), suggesting a protective role of HMGB1 silencing on liver function(26). Histological analysis showed that HMGB1 silencing reduced lipid accumulation in the liver, supporting that HMGB1 contributes to metabolic dysregulation and inflammation in T2D(27). This reduction in lipid deposition and inflammation may be attributed to improved maintenance of liver tissue integrity due to HMGB1 silencing(28). HMGB1 silencing has been shown to modulate upstream signals, such as reducing oxidative stress or inhibiting Toll-like receptor (TLR) activation, which are key drivers of NF-κB activation. Additionally, targeting silencing of upstream mediators of proinflammatory cytokines can attenuate NF-κB signaling(29). Our study pushed the needle forward in emphasizing the inflammatory role of HMGB1 in metabolic diseases, particularly T2D. Previous studies have shown that HMGB1 is a crucial proinflammatory mediator contributing to the pathogenesis of liver injury and insulin resistance(11). Our results align with and expand upon these findings by showing that silencing HMGB1 reduces hepatic inflammation and systemic HMGB1 levels, contributing to better glycemic control(30). The observed reduction in circulating HMGB1 levels and subsequent improvement in glucose homeostasis with exercise are consistent with other reports demonstrating the anti-inflammatory effects of physical activity in metabolic disease models(6). Our findings indicate that the combination of HMGB1 knockdown and exercise produces beneficial effects, enhancing insulin sensitivity and improving metabolic outcomes compared to either one(14). An incidental yet intriguing finding was the differential modulation of metabolite profiles in liver versus skeletal muscle tissues following exercise in T2D mice. The more extensive changes observed in liver metabolites suggest that the liver may be more responsive to HMGB1 silencing and exercise interventions than skeletal muscle. This highlights tissue-specific responses to metabolic regulation in T2D, which could be important for designing more targeted therapies(31). This effect shows interesting metabolic responses between T2D and non-T2D models with HMGB1 knockdown and exercise(11). While exercise improved insulin signaling and glycemic control across both groups, the magnitude of improvement was more pronounced in T2D mice(32). Metabolic cage analyses revealed that HMGB1^Δ^ T2D mice under basal (sedentary) conditions exhibit a respiratory quotient (RQ) close to 0.6, indicating a primary reliance on lipids, proteins, and amino acids for energy(33). In contrast, following exercise, these mice showed an RQ shift toward healthier levels, reflecting enhanced glucose metabolism and a significant metabolic shift from lipid reliance to carbohydrate utilization(34). This shift was accompanied by increased energy expenditure, oxygen consumption, and effective carbohydrate usage, as opposed to the inefficient energy metabolism observed in sedentary T2D controls (Fig. 4). Furthermore, metabolite profiling demonstrated distinct changes in circulating and tissue-specific metabolites post-exercise, with HMGB1^Δ^ T2D mice showing greater modulation of carbohydrate-related metabolites in the liver and skeletal muscle (Fig, 5 and 7). These findings highlight the synergistic effects of HMGB1 knockdown and exercise in driving metabolic flexibility and improved glucose utilization in T2D, contrasting sharply with the metabolic rigidity of sedentary T2D controls(6, 30). This raises interesting questions about the differential impact of exercise and HMGB1silencing in disease versus healthy state. While this difference reflects the inflammatory and metabolic dysregulation in T2D, it also emphasizes the potential of therapeutic benefit by targeting HMGB1 specifically for T2D-associated liver and metabolic dysfunction(35). Related to metabolomic profile analysis post-exercise HMGB1^Δ^ T2D in the liver, our untargeted metabolomic analysis evidenced an increase in key mediators of lipolysis but also enhanced carbohydrate metabolites in HMGB1^Δ^ T2D mice post-exercise. DL-Carnitine and 3-formyl-l-tyr, a derivative of tyrosine, were shown to be increased in HMGB1^Δ^ T2D mice post-exercise (https://pubchem.ncbi.nlm.nih.gov/compound/Tyrosine). 2-amino-3-methyl was also increased in livers from HMGB1^Δ^ T2D mice post-exercise. This is essential for breakdown of tryptophan and vitamin B-6 as an energy source for skeletal muscle (https://pubchem.ncbi.nlm.nih.gov/compound/2-Amino-3-methyl-9H-pyrido_2_3-b_indole).

Enhanced lipid and carbohydrate metabolic intermediates show an enhanced effect in insulin tissues like skeletal muscle as shown in Figure 7. Skeletal muscle untargeted metabolomics showed great benefits from exercise in HMGB1^Δ^ T2D mice post-exercise with major distinct metabolites evidenced between both the liver and skeletal muscle(36–38). The major differences we observed were identified in HMGB1^Δ^ T2D mice post-exercise compared to HMGB1^Δ^ non- T2D mice post-exercise in Figure 7B and 7F(39, 40). We evidenced minimum changes in the amount of decreased or increased metabolites in skeletal muscle between non-T2D and T2D HMGB1^Δ^ mice (Fig.7B). However, we observed an increase in significantly decreased metabolites in skeletal muscle (from 9 to 32) when comparing HMGB1^Δ^ T2D mice undergoing exercise to sedentary (Fig.7F). This points to a systemic effect of liver mediated silencing of HMGB1 enhanced by exercise in T2D that is not as pronounced in non-T2D mice. Therefore, we hypothesize that not only local silencing in hepatic HMGB1 increases insulin sensitivity increases in our model but that decreasing circulating HMGB1 might have a high effect in other insulin sensitive tissues like skeletal muscle and therefore increasing the highly beneficial effects we observe locally in the liver, systemically in peripheral tissues insulin sensitivity and the enhanced metabolic shift to carbohydrate use systemically enhanced by exercise due to the involvement of the skeletal muscle as evidenced by our supporting data in Supplemental Figures 1 and 2. We decided to perform in vitro skeletal muscle contraction model (EPS) confirmed that the beneficial effects of HMGB1 knockdown are enhanced when combined with exercise that extends to skeletal muscle, further distinguishing our work from prior studies(41). Finally, another unexpected result was the improved bone mineral density in HMGB1^Δ^ T2D mice but exponentiated post exercise, suggesting that HMGB1 might also regulate bone reabsorption and density under diabetic conditions(42). We speculate that the metabolic improvements observed in HMGB1^Δ^ T2D mice post-exercise enhance carbohydrate metabolism and influence bone metabolism, potentially through mechanisms tied to osteoblast and osteoclast activity. HMGB1 protein has been shown to regulate osteoclastogenesis via direct actions on osteocytes and osteoclasts in vitro(43). In our model, reducing HMGB1 in combination with exercise could indirectly suppress excessive osteoclast activity. At the same time, the enhanced energy metabolism, evidenced by increased carbohydrate utilization and respiratory quotient (Fig. 4), may support osteoblastic activity and bone formation. This metabolic shift aligns with findings from exercise studies showing improved bone density in diabetes through energy metabolism pathways(44, 45). Our results in Fig. 6 demonstrate increased bone mineral density and lean muscle mass in HMGB1^Δ^ T2D mice post-exercise, likely driven by enhanced systemic energy availability and metabolic flexibility. This suggests that exercise induced shifts in glucose metabolism and energy expenditure improve glycemic control and positively influence skeletal health in diabetes, potentially through combined effects on insulin signaling, osteoblast activation, and reduced HMGB1 mediated osteoclastogenesis(46). We next need to discuss the limitations of our studies. There is an emergent need for a comprehensive assessment of both hepatic and systemic metabolic changes in response to targeted HMGB1 interventions(6). We addressed this scientific limitation by employing metabolomic profiling in plasma and liver tissues and analyzing exercise effects in different metabolic compartments, including skeletal muscle. This integrated approach allowed us to capture the broad impacts of HMGB1 knockdown, providing new insights into how localized hepatic interventions can influence systemic metabolism and inflammatory responses. Another limitation is the use of only male mice, which does not allow us to evaluate potential sex-specific differences in response to HMGB1 silencing and exercise interventions. But the lack of females in this experiment is given to minimize variability associated with hormonal cycles in females(47). However, it overlooks important sex-based physiological differences. Notably, female mice exhibit greater resistance to streptozotocin (STZ)-induced diabetes compared to males, likely due to protective effects of estrogen on glucose metabolism(6, 48). Additionally, while the exercise regimen employed was adequate, the lack of varying exercise intensities and modalities limits the translational and generalizability of our findings to other types of physical activity. Future studies will target how different forms of exercise might differentially impact metabolic outcomes in T2D.

## 5. CONCLUSIONS

In conclusion, our studies demonstrate that liver HMGB1 silencing in T2D mice undergoing exercise induces carbohydrate metabolism, decreased hyperglycemia and liver protection. Our results propose a novel strategy to mitigate liver damage, improve metabolic health, and enhance glycemic control in T2D that can be translated to clinical applications. Future directions will be aimed at further investigating the Akt and FoxO1 directed mechanism that our group has evidenced in the role of HMGB1 in enhancing insulin sensitivity to understand its contribution to systemic inflammation and metabolic dysfunction. Furthermore, clinical translation of these findings will require the development of exercise-driven HMGB1-targeting approaches that are safe and effective in humans, as well as the exploration of personalized exercise prescriptions to maximize therapeutic outcomes for individuals with T2D.

## AVAILABILITY OF DATA AND MATERIALS

The data that supports the findings of this is available in the MENDELEY DATA repository and will be given access upon reasonable request: Mota Alvidrez, Roberto (2024), “Exercise Training Enhances Insulin Sensitivity with Metabolic and Glycemic Control Driven by Hepatic Silencing of HMGB1 in Type 2 Diabetic Mice”, Mendeley Data.

## AUTHOR CONTRIBUTIONS

GMB: Writing – review & editing, Writing – original draft, Software, Methodology, Investigation, Formal analysis, Data curation. QJ, PP, GS, HB, EB: Validation, Resources, Methodology, Formal analysis, Data curation. JC, HC: Validation, Resources, Formal analysis, Data curation. FCM: Visualization, Validation, Supervision, Conceptualization, Data curation. RIMA: Writing – review & editing, Writing – original draft, Visualization, Validation, Supervision, Software, Resources, Project administration, Methodology, Investigation, Funding acquisition, Formal analysis, Data curation, Conceptualization. All authors contributed to editorial changes in the manuscript. All authors read and approved the final manuscript.

## ETHICS APPROVAL AND CONSENT TO PARTICIPATE

The ethics committee of the University of New Mexico (UNM) Institutional Animal Care and Use Committee (IACUC) and Animal Welfare Committee approved all animal experiments. All live animal studies were conducted ethically, following relevant guidelines and regulations at the University of New Mexico under protocol number 23-201405-HSC.

## ACKNOWLEDGMENT

We would like to thank Dr. Matthew Neal and Dr. Timothy Billiar at the University of Pittsburgh for providing the HMGB1 flox mice that were used in our studies. We would like to thank Dr. Michael Deyhle and his group for preparing C2C12 cells, sharing EPS protocols, counseling and guidance with interpretation of our in vitro results. We would like to express our gratitude to Dr. Meilian Liu for her expert guidance in DEXA Scan imaging and metabolic cage studies. We would like to thank Jinhua Chi and Haiwei Gu for processing, analysis and interpretation of all our untargeted metabolomics. Additionally, we appreciate the support provided by Dr. Mathew Campen and his laboratory in the development of metabolomic assays and the analysis of metabolomic data.

## FUNDING

This research was funded by NHLBI R25HL145817 to RIMA, NCATS KL2 TR001448 funding for RIMA (UNM HSC CTSC).

## CONFLICT OF INTEREST

The authors declare no conflict of interest.

---

**Supplementary Figure 1.**
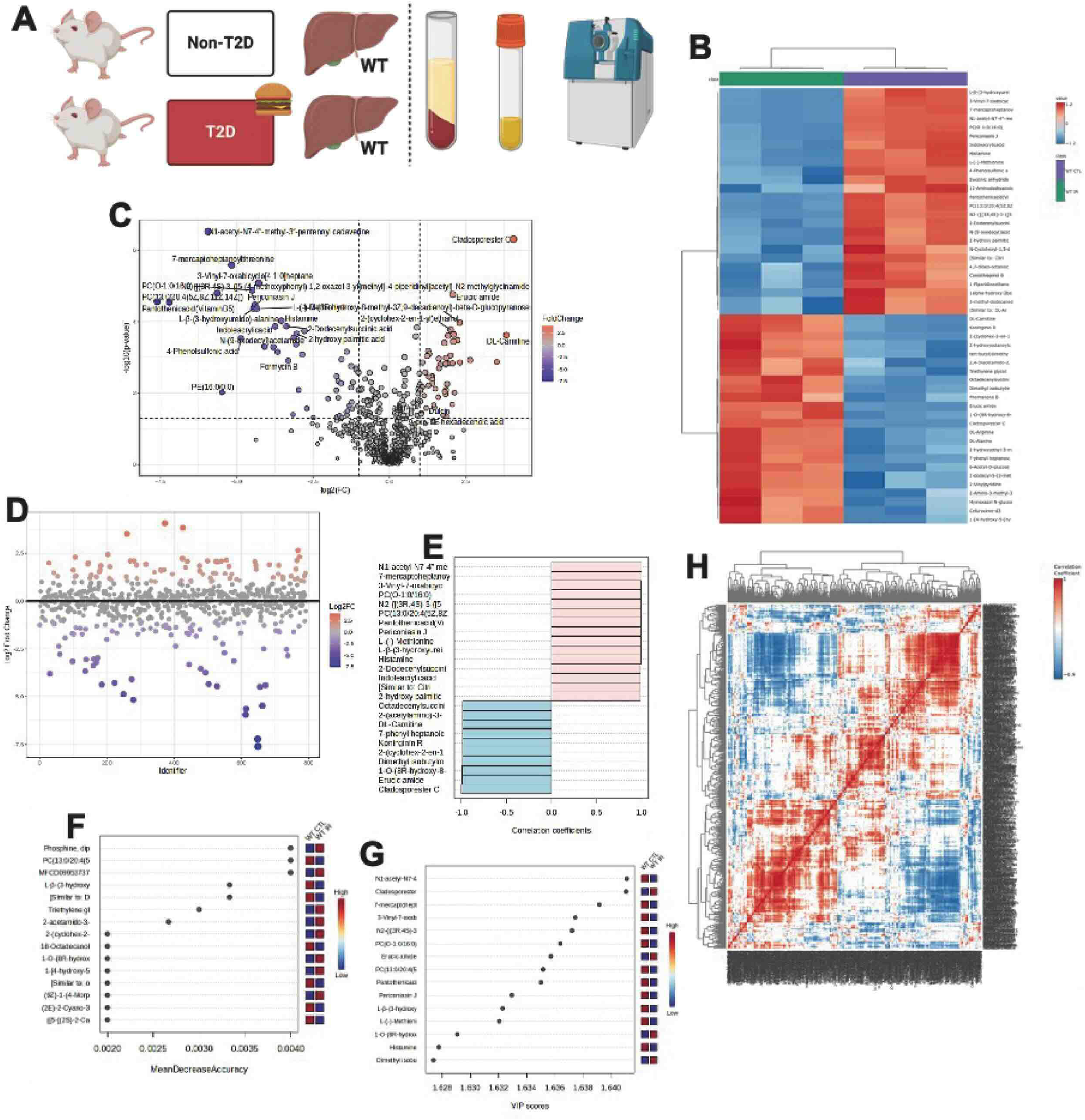

**Supplementary Figure 2.**
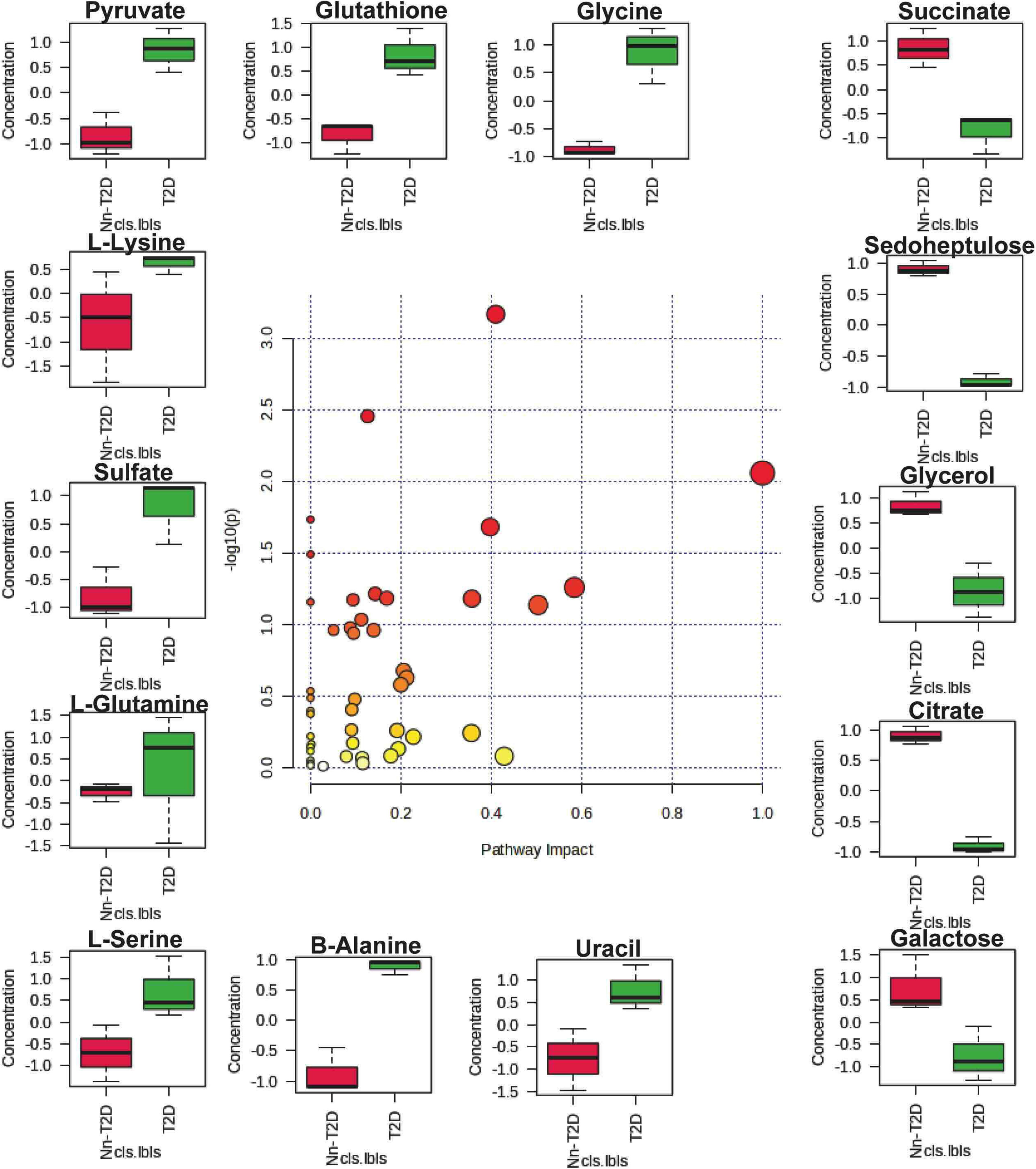

**Supplementary Figure 3.**
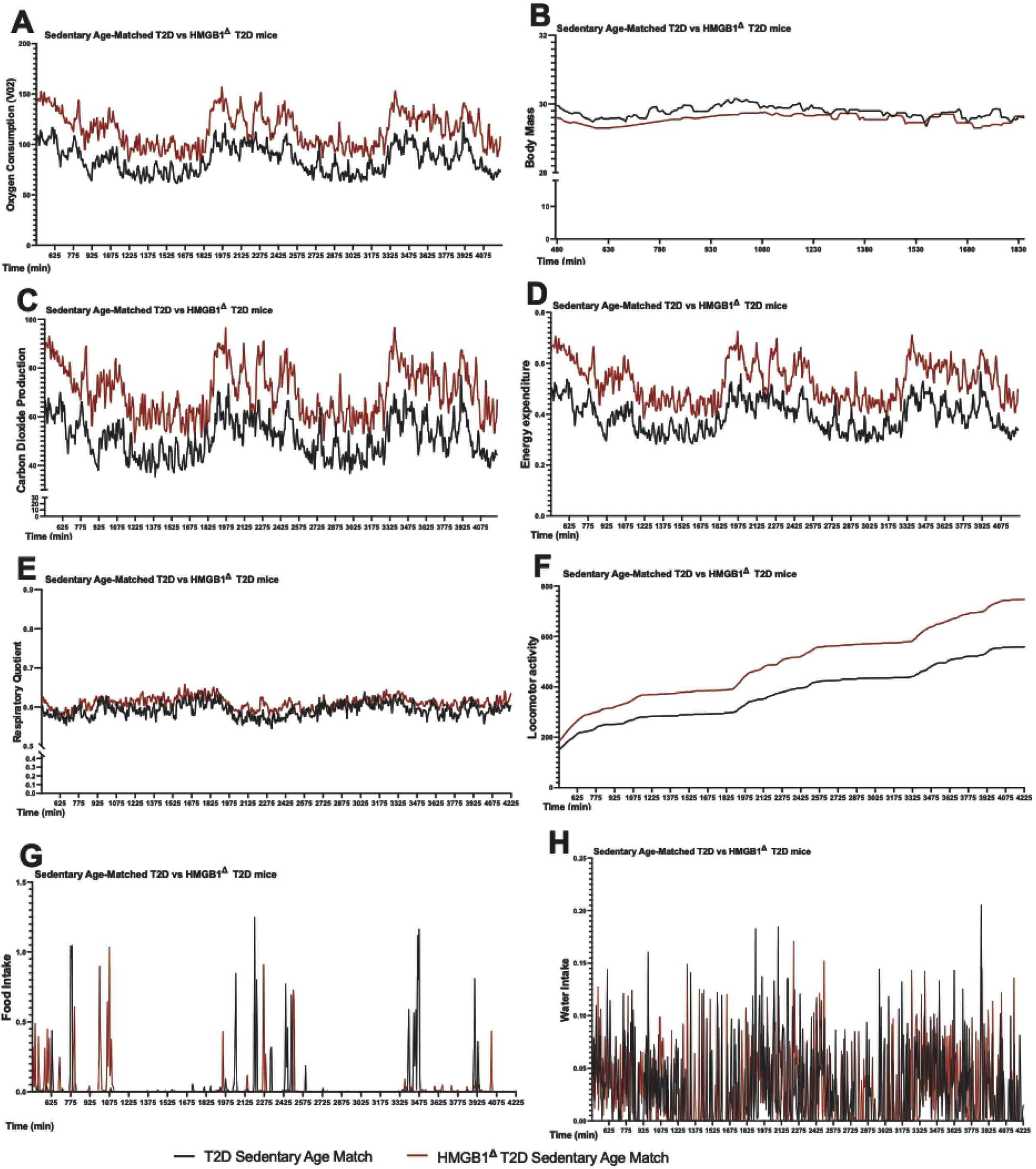

**Supplementary Figure 4.**
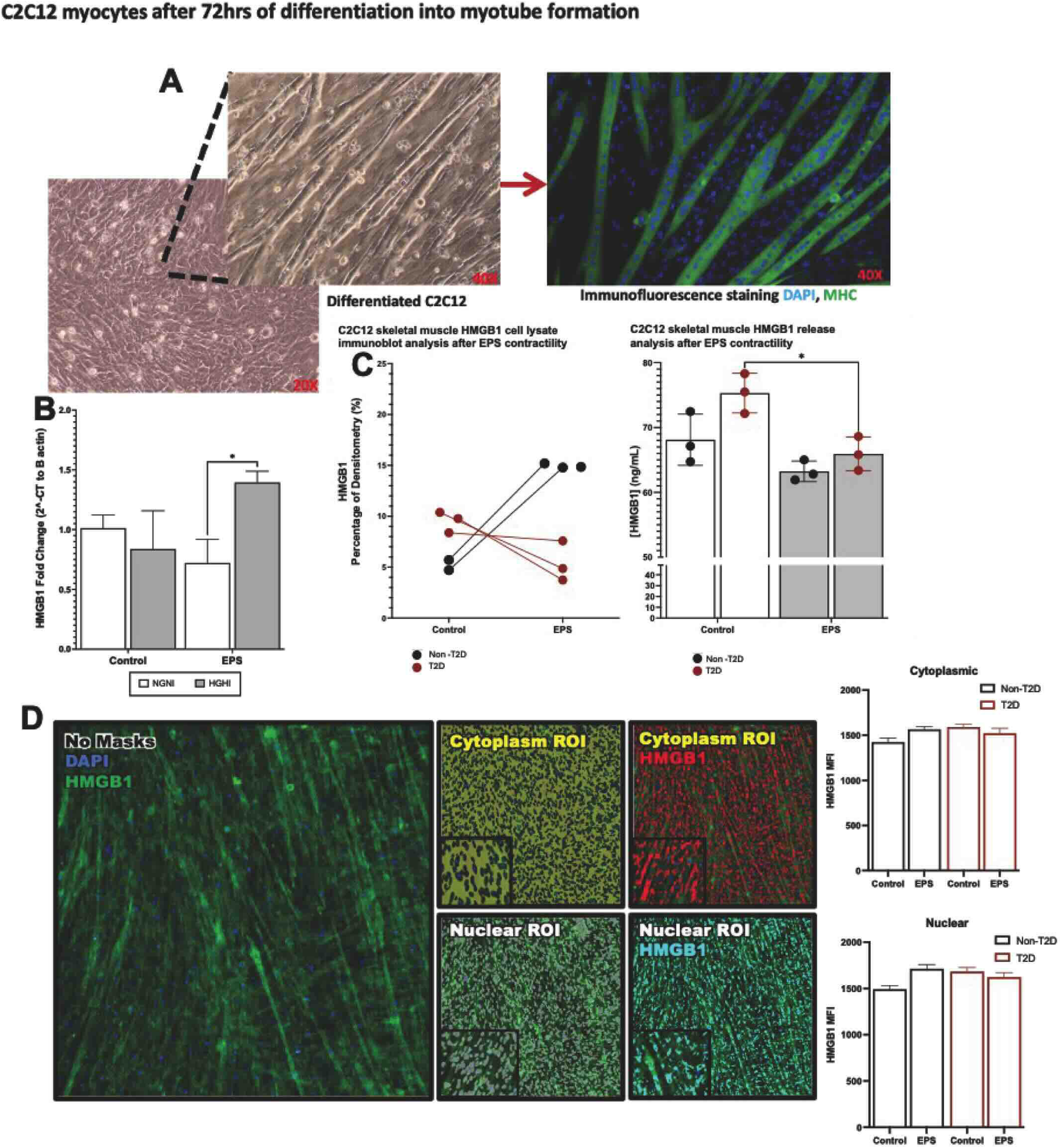

## Notes

### Competing Interest Statement

The authors have declared no competing interest.

