## Supplemental Material for "Exercise Conditioning Enhances Insulin Sensitivity with Metabolic and Glycemic Control Driven by Hepatic Silencing of HMGB1 in Type 2 Diabetic Mice"

### Supplemental Methods

*Metabolic cage analysis*

To gather information from metabolic cages, each mouse was housed individually in a specialized metabolic cage designed to maintain a controlled environment throughout the experiment. This setup enabled precise measurements of key metabolic parameters, including energy expenditure, respiratory exchange ratio (RER), and physical activity levels. An advanced animal monitoring system facilitated real-time data collection and analysis of the mice's metabolic functions. The cages were carefully controlled to ensure the well-being of the mice and the accuracy of the measurements. The ambient temperature was kept constant at 23°C, providing optimal physiological comfort. The lighting system was also programmed to follow a 12-hour light/dark cycle, mimicking natural conditions. This setup minimized stress and allowed for the assessment of circadian physiological patterns.

*Liver and plasma untargeted metabolomic complete methods*

For tissue preparation, approximately 20 mg of each sample was homogenized in 200 µL of MeOH:PBS (4:1, v:v, containing 1,810.5 µM $^{13}C_3$-lactate and 142 µM $^{13}C_5$-glutamic acid) in an Eppendorf tube using a Bullet Blender homogenizer (Next Advance, Averill Park, NY). An additional 800 µL of the same MeOH:PBS solution was added and vortexed for 10 seconds. Samples were stored at -20$^{o}$ C for 30 min and then sonicated in an ice bath for 30 minutes. Centrifugation at 14,000 RPM for 10 min at 4$^{o}$ C was performed, and 800 µL of the supernatant was transferred to a new Eppendorf tube. The samples were dried under vacuum using a CentriVap Concentrator (Labconco, Fort Scott, KS). Before MS analysis, the residues were reconstituted in 150 µL of 40% PBS/60% ACN. A pooled quality control (QC) sample was prepared by combining aliquots from all study samples. All LC-MS experiments were conducted on a Thermo Vanquish UPLC-Exploris 240 Orbitrap MS instrument (Waltham, MA). Each sample was injected twice: 10 μL for analysis in negative ionization mode and 4 μL for positive ionization mode. Chromatographic separations were performed using hydrophilic interaction chromatography (HILIC) on a Waters XBridge BEH Amide column (150 x 2.1 mm, 2.5 μm particle size, Waters Corporation, Milford, MA). The flow rate was 0.3 mL/min, with the autosampler maintained at $4^{o}$ C and the column compartment at $40^{o}$ C. The mobile phase consisted of Solvent A (10 mM ammonium acetate, 10 mM ammonium hydroxide in 95% $H_2O$/5% ACN) and Solvent B (10 mM ammonium acetate, 10 mM ammonium hydroxide in 95% ACN/5% $H_2O$). After an initial 1-minute isocratic elution with 90% Solvent B, the percentage of Solvent B decreased to 40% over 11 minutes, held for 4 minutes, and then returned to 90% to prepare for the next injection. Mass spectrometric data were collected over 70 to 1050 m/z using an electrospray ionization (ESI) source.

Peak identification in the MS spectra utilized an extensive in-house chemical standards library (~600 aqueous metabolites) and comparisons with external databases, including the Human Metabolome Database (HMDB), LipidMaps, METLIN, mzCloud, Metabolika, and ChemSpider. MS data extraction employed an absolute intensity threshold of 1,000, with a mass accuracy limit of 5 ppm. Identification and annotation leveraged retention time (RT), exact mass (MS), MS/MS fragmentation patterns, and isotopic distribution data. Data processing for aqueous metabolomics was conducted using Thermo Compound Discoverer 3.3 software, which handled peak picking, alignment, and normalization. To ensure rigor only signals with a coefficient of variation (CV) < 20% across QC pools and those detected in >80% of samples were included for further analysis.

*Tissue preparation*

Whole blood was collected and transferred directly to EDTA2 plasma tubes, which were centrifuged for plasma separation. Plasma was analyzed using HMGB1 Elisa (MyBioSource, MBS701378). The liver was equally split for the different studies. Part of the liver was homogenized; total protein was extracted for immunoblot and RNA was isolated for PCR analysis as previously described. To assess liver function, plasma samples collected previously were diluted at a ratio of 1:10 to prepare for the measurements of the enzyme Aspartate aminotransferase (AST). These measurements were conducted using the DRI-CHEM analyzer by Heska (Loveland, CO). The procedure was performed based on the manufacturer's kit guidelines and recommended protocol to ensure the accuracy and reliability of the results. For the histological staining, part of the livers was immersed and fixed in 4 % PFA for 4 h, followed by overnight incubation in a 30 % sucrose solution for cryoprotection. Livers were embedded in OCT solution and cryosectioned into 10 µm-thick sections. According to the manufacturer's protocol, Hematoxylin and eosin (H&E) staining (Abcam, ab245880) was performed on the same tissue sections. Bright field microscope images were captured for analysis at 20X imaging focus. Liver, serum/plasma, and skeletal muscle were sent to Arizona State University for untargeted metabolic analysis, described in the following section.

*Simulated Muscle Contraction via Electrical Pulse Stimulation in C2C12 cells*

C2C12 myoblasts (American Type Culture Collection, Cat. No. CRL-1772, RRID: CVCL_0188, Manassas, VA) were seeded in 6-well cell culture dishes at 5 x $10^3$ cells/$cm^2$. This protocol has been used previously [20] and has been shown to recapitulate muscle adaptations and responses to contractions that occur in vivo, such as hypertrophy [21]. Cells were cultured at 37°C and 5% $CO_2$ in growth medium (Dulbecco's Modified Eagle's Medium [DMEM] +10% fetal bovine serum and 1% penicillin/streptomycin) until 90% confluence was reached. The medium was then switched to low-serum medium (DMEM + 2% horse serum and 1% penicillin/streptomycin) to encourage myotube formation. After 4 days, myotubes were formed and were ready for experimentation. To induce contractions, myotubes were exposed to electrical pulse stimulation (EPS) using an electrical pulse multichannel culture pacer (C-Pace 100, IonOptix, Dublin, Ireland). EPS pulses were 12 V, 2 ms pulses in duration at a frequency of 1 Hz. EPS was applied for 8 hours.

### Supplemental Figures and Legends

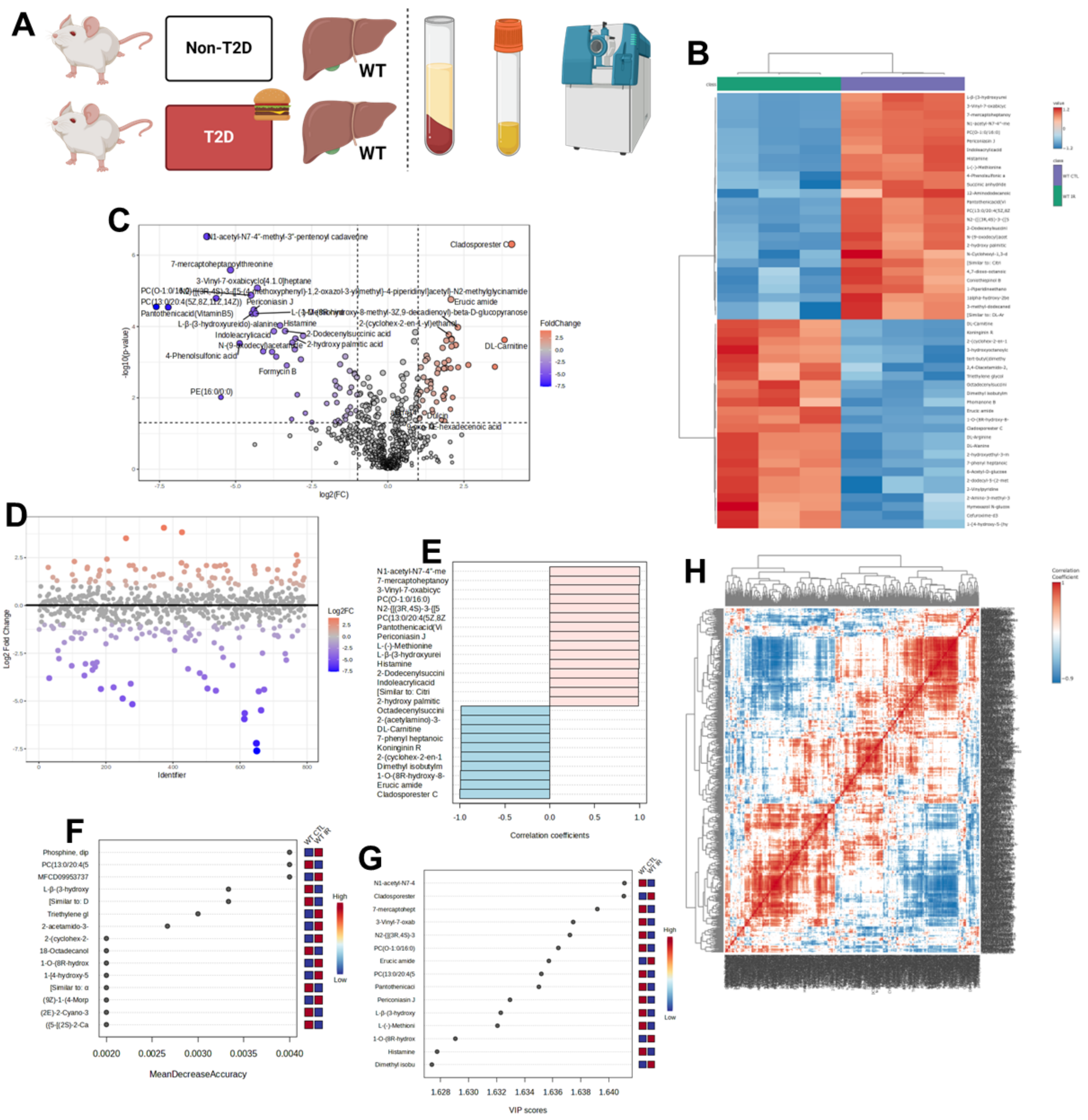


**Supplementary Figure 1. Circulating metabolite analysis identified overriding clustered differences driven by T2D in sedentary control mice**. A) Diagram of the different mouse models used in this study post-exercise HMGB1$^{\Delta}$ non-T2D and HMGB1$^{\Delta}$ T2D, followed by the procedure performed, B) Heat map displays the up- and down-regulated metabolites in the plasma, C) Volcano plot from plasma metabolites showing the different amount of significant up, down, and insignificant metabolite changes, D) Fold change (FC) in metabolites levels in plasma samples, E) OPLS plot, F) SPLS plot, G) VIP score plot, H) correlation heat map. All data is represented as mean±SEM. N of 4 for non-T2D exercise, 5 for T2D.

### Supplemental Figure 2

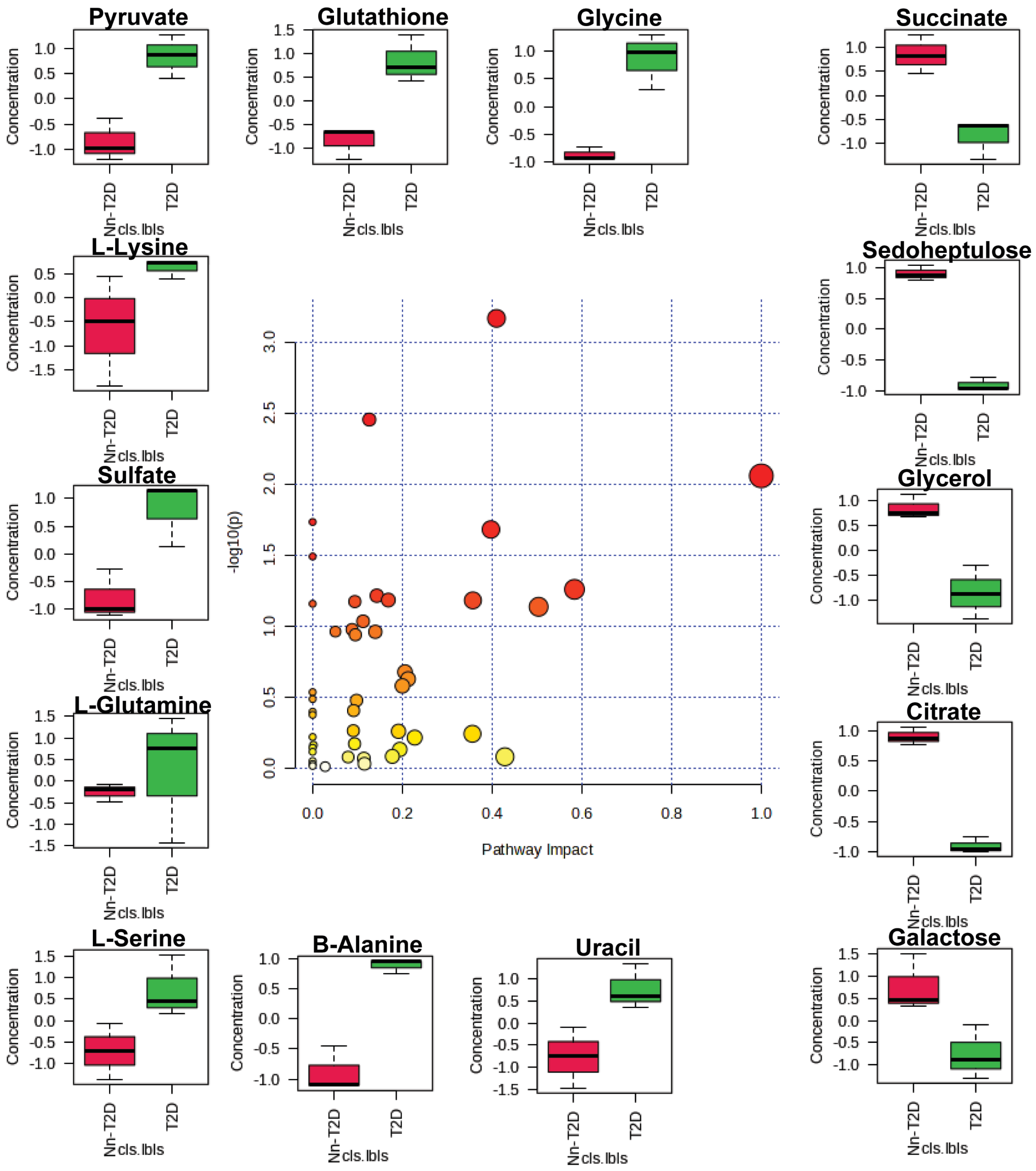

**Supplementary Figure 2. Circulating plasma metabolite pathway impact comparison from sedentary non-T2D vs T2D.** All data is represented as mean±SEM. N of 4 for non-T2D exercise, 5 for T2D.

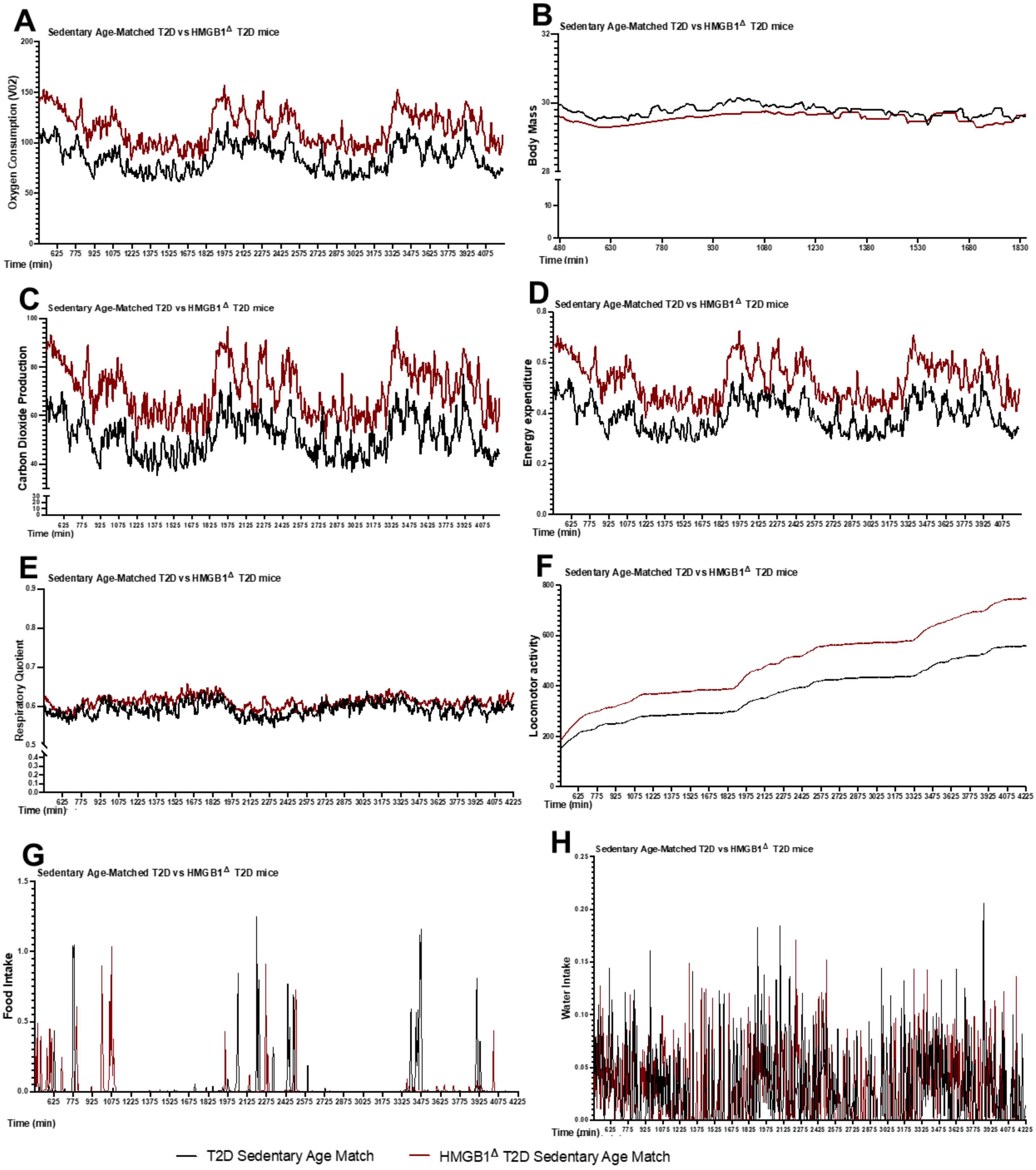
A
Sedentary Age-Matched T2D vs HMGB1Δ T2D mice
Oxygen Consumption (V02)
Time (min)
B
Sedentary Age-Matched T2D vs HMGB1Δ T2D mice
Body Mass
Time (min)
C
Sedentary Age-Matched T2D vs HMGB1Δ T2D mice
Carbon Dioxide Production
Time (min)
D
Sedentary Age-Matched T2D vs HMGB1Δ T2D mice
Energy expenditure
Time (min)
E
Sedentary Age-Matched T2D vs HMGB1Δ T2D mice
Respiratory Quotient
Time (min)
F
Sedentary Age-Matched T2D vs HMGB1Δ T2D mice
Locomotor activity
Time (min)
G
Sedentary Age-Matched T2D vs HMGB1Δ T2D mice
Food Intake
Time (min)
H
Sedentary Age-Matched T2D vs HMGB1Δ T2D mice
Water Intake
Time (min)
T2D Sedentary Age Match
HMGB1Δ T2D Sedentary Age Match

**Supplementary Figure 3. Metabolic cage analysis of control sedentary age matched HMGB1$^{\Delta}$ T2D vs T2D.** A) Linear graph representing the difference in oxygen consumption age match of sedentary T2D, HMGB1Δ T2D mice, B) Linear graph representing the difference in body mass of age match mice, C) Linear graph representing the difference in Carbon dioxide production of age match mice, D) Linear graph representing the difference in Energy Expenditure of age match mice, E) Linear graph representing the difference in Respiratory rate of age match mice, F) Linear graph representing the difference in Locomotor activity of age match mice, G) Linear graph representing the difference in Food intake of age match mice, H) Linear graph representing the difference in Water Intake of age match mice. All data is represented as mean±SEM. N of 4 for T2D, and 4 for HMGB1$^{\Delta}$ T2D mice.

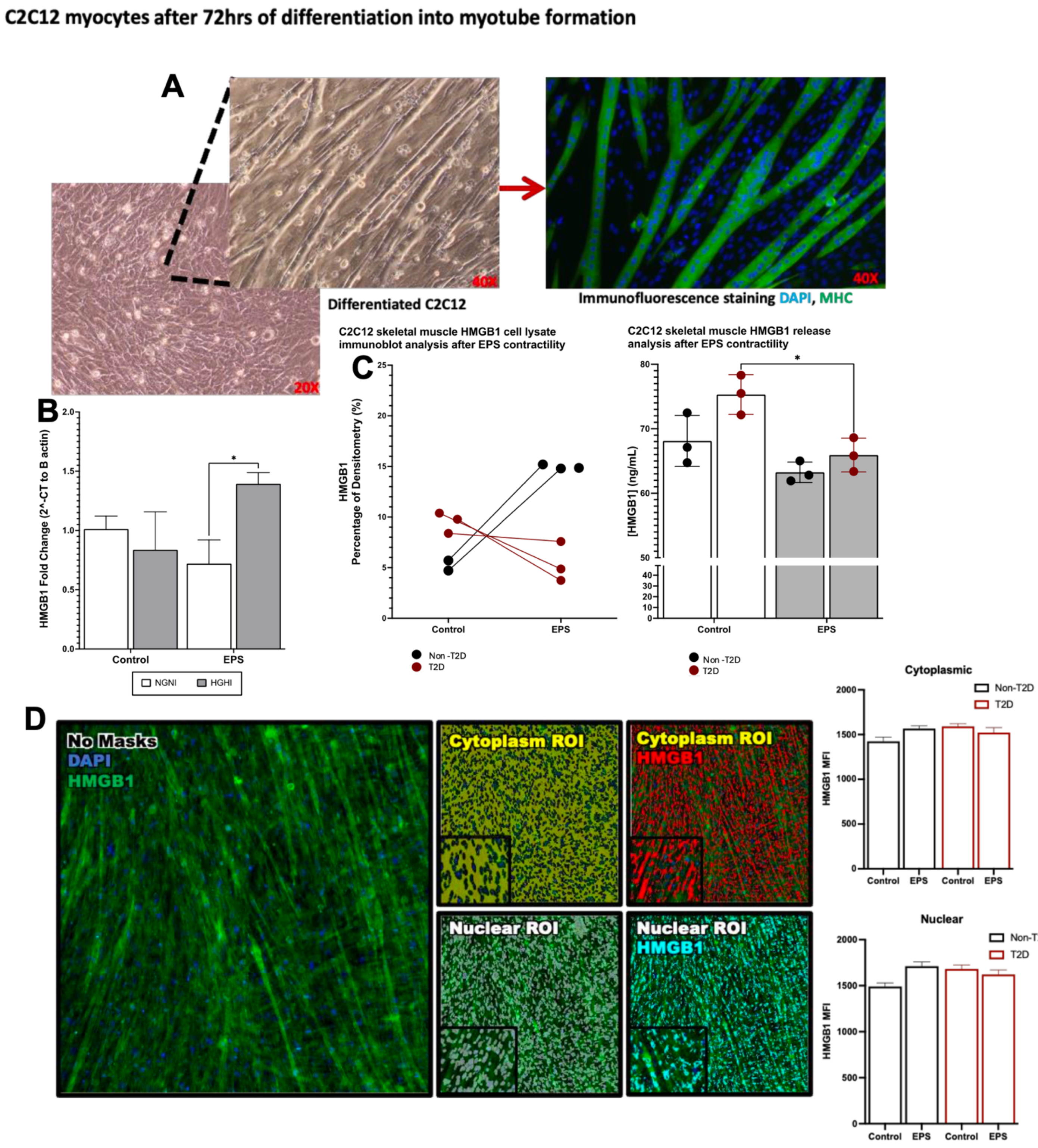


**Supplemental Figure 4. Exercise mimicking stimulation in vitro with EPS evidence inhibition of HMGB1 cellular release in skeletal muscle cells.** A) Imaging of differentiated c2c12 cells, immunofluorescence detecting HMGB1, B) qRT-PCR analysis of HMGB1 gene expression in c2c12 cells with and without EPS, C) Cell lysate immunoblot analysis after EPS fror HMGB1 and HMGB1 release analysis, C) Cellomics of differentiated c2c12 cells showing HMGB1 in cytoplasm and nuclear with bar graphs representing the results. All data is represented as mean ±SEM. N of 4 for T2D, and 4 for HMGB1$^{\Delta}$ T2D mice.
